# An Integrative Approach to Dissect the Drug Resistance Mechanism of the H172Y Mutation of SARS-CoV-2 Main Protease

**DOI:** 10.1101/2022.07.31.502215

**Authors:** Joseph Clayton, Vinicius Martins de Oliveira, Mohamed Fouad Ibraham, Xinyuanyuan Sun, Paween Mahinthichaichan, Mingzhe Shen, Rolf Hilgenfeld, Jana Shen

## Abstract

Nirmatrelvir is an orally available inhibitor of SARS-CoV-2 main protease (Mpro) and the main ingredient of PAXLOVID, a drug approved by FDA for high-risk COVID-19 patients. Recently, a rare natural mutation, H172Y, was found to significantly reduce nirmatrelvir’s inhibitory activity. As the COVID-19 cases skyrocket in China and the selective pressure of antiviral therapy builds up in the US, there is an urgent need to characterize and understand how the H172Y mutation confers drug resistance. Here we investigated the H172Y Mpro’s conformational dynamics, folding stability, catalytic efficiency, and inhibitory activity using all-atom constant pH and fixed-charge molecular dynamics simulations, alchemical and empirical free energy calculations, artificial neural networks, and biochemical experiments. Our data suggests that the mutation significantly weakens the S1 pocket interactions with the N-terminus and perturbs the conformation of the oxyanion loop, leading to a decrease in the thermal stability and catalytic efficiency. Importantly, the perturbed S1 pocket dynamics weakens the nirmatrelvir binding in the P1 position, which explains the decreased inhibitory activity of nirmatrelvir. Our work demonstrates the predictive power of the combined simulation and artificial intelligence approaches, and together with biochemical experiments they can be used to actively surveil continually emerging mutations of SARS-CoV-2 Mpro and assist the discovery of new antiviral drugs. The presented workflow can be applicable to characterize mutation effects on any protein drug targets.

## Introduction

The COVID-19 pandemic is still ongoing and remains a major global health threat. At the end of 2021, the U.S. Food and Drug Administration (FDA) issued an Emergency Use Authorization for Pfizer’s PAXLOVID to treat mild-to-moderate COVID-19 cases^1,2^. In a recent clinical trial for high-risk non-hospitalized adults with COVID-19^3^, PAXLOVID reduced the risk of progression to severe disease by 89% as compared to placebo. This antiviral drug is a ritonavir-boosted formulation of nirmatrelvir (PF-07321332), an orally available inhibitor of the SARS-CoV-2 main protease (Mpro). Mpro, which is also known as 3CLpro or Nsp5, is a cysteine protease essential to the viral replication process as it cleaves the majority of the polyproteins pp1a and pp1ab into nonstructural proteins which form a part of the viral replication complex^4^. Nirmatrelvir is a reversible covalent peptidomimetic inhibitor, which binds to the active site of Mpro and inhibits its proteolytic activity^5^. Although Mpro is one of the most conserved proteins among coronaviruses^4^, the rapid and constant evolution of the viral genome raises great concern of potential emergence of antiviral resistance. Several biochemical studies, however, showed that the prevalent Mpro mutants in the Variants of Concern or Variants of Interest declared by the World Health Organization (WHO), such as G15S (Lambda), K90R (Beta), and P132H (Omicron), are still susceptible to nirmatrelvir, with IC_50_ values and catalytic efficiencies similar to the wild type (WT) Mpro^6–8^. Nevertheless, biochemical assays of several infrequent natural substitutions, e.g., H164N, H172Y, and Q189K, are associated with reduced activities of nirmatrelvir, among which H172Y caused the largest reduction in the inhibitory activity, with a 233-fold increase in the *Ki* value of nirmatrelvir according to a disclosure by Pfizer^1^. Although H172Y is a rare mutation (found in only a few entries of the database GISAID^9^), it may become favored in the future under the selection pressure of nirmatrelvir therapy. Thus, understanding the antiviral resistance mechanism is important and urgently needed.

Motivated by the aforementioned need, we investigated the effect of the H172Y mutation on Mpro’s structure, stability, and binding with nirmatrelvir using a battery of state-of-the-art computational approaches, including the all-atom constant pH and fixed-charge molecular dynamics (MD), alchemical free energy simulations, empirical folding and binding free energy calculations, and artificial neural networks. As an experimental structure of the H172Y Mpro was unavailable at the start of the study, the computational work was solely based on an in silico mutated structure model. The simulations revealed that the H172Y substitution disrupts the S1 pocket interactions with the N-terminus of the opposite protomer and perturbs the conformation of the oxyanion loop. The empirical calculations predicted a decreased stability for the H172Y Mpro. The empirical and alchemical free energy simulations predicted a decreased binding affinity with nirmatrelvir. These results were verified with the thermal stability, enzyme kinetics, and inhibitory activity measurements. The MD data also corroborate with the newly reported X-ray structure models of H172Y Mpro^10^.

## Results and Discussion

### Molecular dynamics simulations of the free and nirmatrelvir-bound H172Y Mpro

We first built a structure model of H172Y Mpro based on the X-ray crystal structure of WT Mpro in complex with nirmatrelvir (PDB id 7vh8, resolution 1.58 A, Fig. 1)^5^ using Modeller. The modeled H172Y Mpro structure is nearly superimposable with the WT, except for a slight displacement of the backbone of Phe140, resulting in a 0.3 Å larger distance between the backbone carbonyl oxygen of Phe140 and the amino nitrogen of Ser1* (asterisk indicates the opposite protomer). The S1 pocket-Ser1* remain intact as in the WT-Mpro (Fig. 1). The protonation states of H172Y Mpro were determined using the generalized Born (GBNeck2) continuous constant pH molecular dynamics (CpHMD) titration simulations^12,13^ with the asynchronous pH replica-exchange protocol for enhanced sampling. The estimated *pK*_a_’s are similar to those of the WT Mpro^15^ and the protonation states at pH 7.5 remain the same (Table S1). Note, consistent with other MD studies^16^ our previous work showed that His172 in the WT Mpro is predominantly neutral at physiological pH, and a switch to the charged state at low pH results in a partial collapse of the S1 pocket^15^; such pH-dependent behavior is removed by the Tyr172 substitution in the mutant.

**Figure 1.**
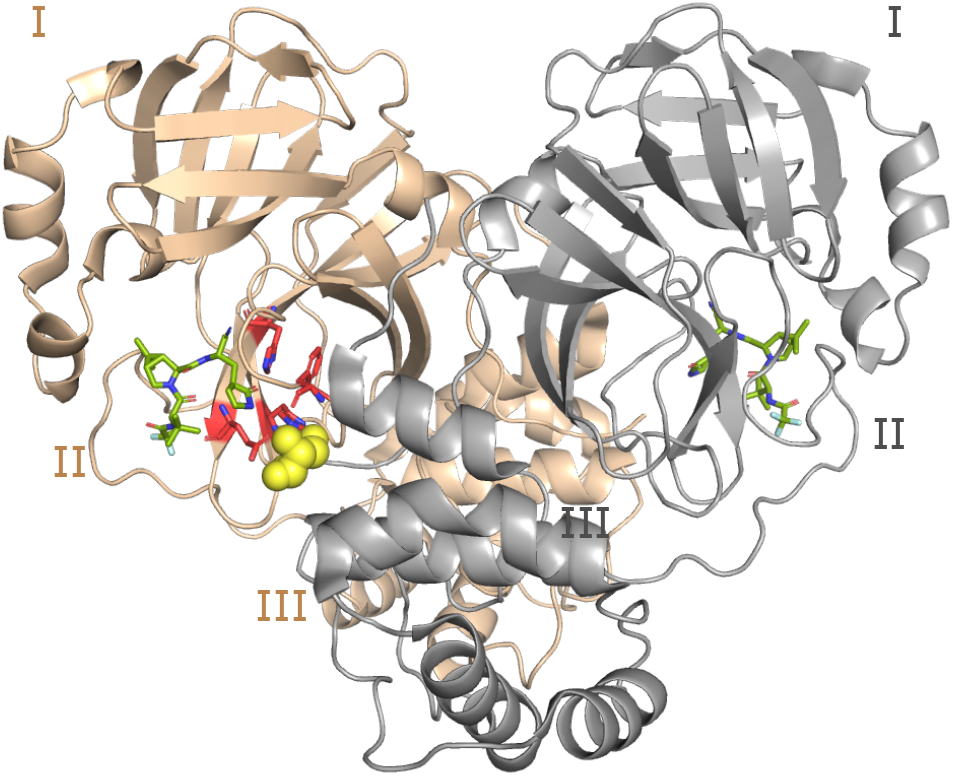
Structure of the WT SARS-CoV-2 Mpro dimer. Cartoon representation of the Mpro dimer bound to nirmatrelvir (PDB ID 7vh8^5^) with protomer A in tan and B (front of the image) in grey. The three domains (I, II, and III) are labeled for each protomer. The S1 pocket residues (Phe140, His163, Glu166, and His172) of protomer A (highlighted in red and shown as sticks) interacts with Ser1^*^ from protomer B (shown in the van der Waals sphere representation). Ser1^*^ forms either a hydrogen bond or salt bridge with Phe140, Glu166, and His173, while His163 forms aromatic stacking with Phe140. The inhibitor nirmatrelvir is shown in green.

Starting from the computationally mutated structure and with the CpHMD determined protonation states, we carried out fixed-charge MD simulations of the free as well as the nirmatrelvir-bound H172Y Mpros using the Amber20 program^17^. As a control, the free and ligand-bound WT Mpros were also simulated starting from the same template structure (PDB id 7vh8) ^5^ and with the same settings. A total of 10 simulations runs were conducted, including 3 trajectories for the free WT/H172Y Mpros and 2 trajectories for the ligand-bound WT/H172Y Mpros, with each trajectory lasting 2 μs (Table S2). In all these trajectories, the overall structure of the Mpro was stable and the inhibitor remained bound (Fig. S1-S2). At the end of our study, one free H172Y Mpro trajectory was also obtained starting from our unpublished X-ray structure of H172Y Mpro (see later discussion).

### S1 pocket interactions with N-terminus* are destabilized in the simulations of free H172Y Mpro

A unique feature of the SARS-CoV/SARS-CoV-2 Mpros is the interactions between the S1 pocket residues and the N-finger (residues 1-9) of the opposite protomer (Fig. 1); these interactions are believed to support the stability of the active site and the Mpro dimerization^4,18^. In particular, three absolutely conserved residues in the S1 pocket, Phe140, Glu166, and His172 form either hydrogen bond (H-bond) or salt bridge with the N-terminus of the opposite protomer (i.e., the backbone of Ser1*, asterisk denotes the opposite protomer), according to the X-ray structures^4,18^ and the previous ^15^ as well as the current WT Mpro simulations (Fig. 2a and Fig. S3). We first consider the H-bond between the backbone carbonyl group of Phe140 and Ser1*. This H-bond remained stable in all the WT Mpro simulations; in contrast, it became disrupted for both protomers after 1 μs in run 1 and almost immediately disrupted in run 2 and run 3 of the H172Y Mpro (Fig. 2b and Fig. S4-S6). The WT Mpro simulations showed that the charged Glu166 and the terminal amine of Ser1^*^ form either a H-bond/salt bridge or electrostatic interaction; in contrast, the Glu166-Ser1^*^ interaction became disrupted for both protomers after 1 μs in run 1 and almost immediately disrupted run 2 and run 3 of the H172Y Mpro (Fig. 2d and Fig. S4-S6).

**Figure 2.**
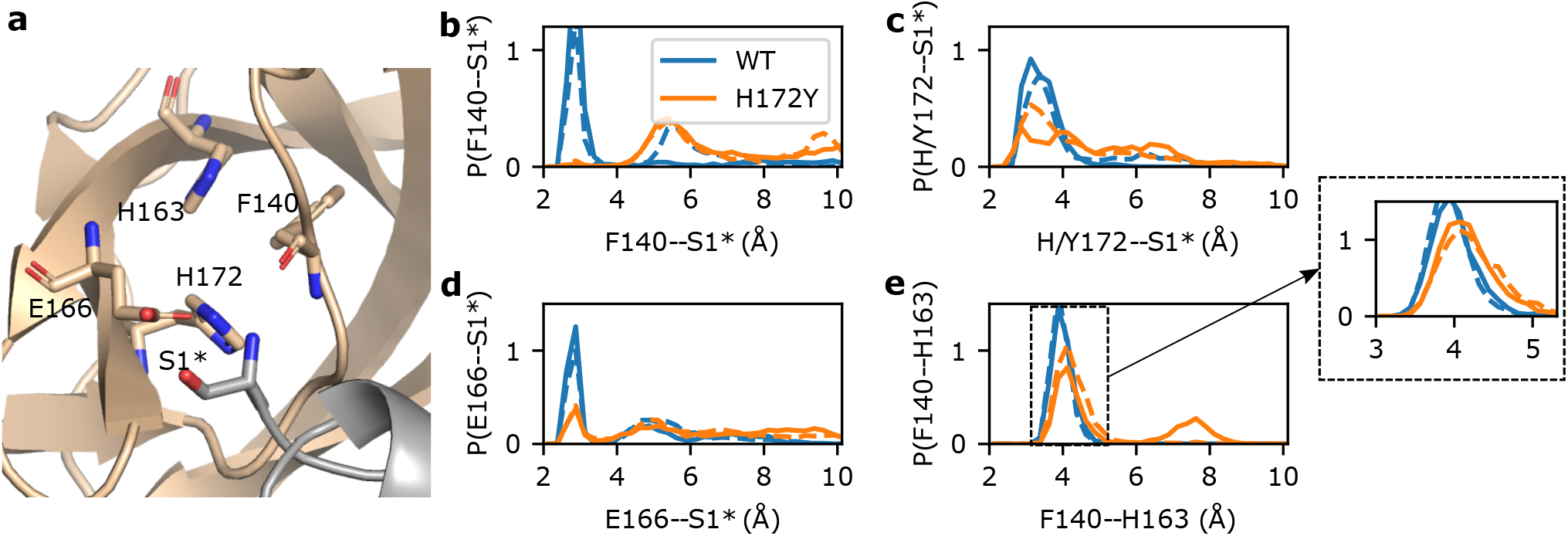
N-terminus interactions with the S1 pocket are destabilized in the simulations of the free H172Y Mpro. **a.** Visualization of the interactions between the S1 pocket and N-terminus* (Ser1*) of the opposite protomer in the WT Mpro. Phe140, His163, Glu166, His172, and Ser1* are shown as sticks. **b,c,d.** Probability distributions of distances between Phe140 (b), Glu166 (c), or His/Tyr172 (d) and Ser1* from the WT and H172Y Mpro simulations. For each Mpro, all three trajectories were used with the first iμs of each trajectory discarded. In b and c, distance was calculated from the N-terminal nitrogen of Ser1* to the backbone carbonyl oxygen of Phe140 (b) or the nearest carboxylate oxygen of Glu166 (c). In d, distance was calculated from the backbone carbonyl oxygen of Ser1* to the nearest imidazole nitrogen of His172 (WT) or from the N-terminal nitrogen of Ser1* to the hydroxyl oxygen of Tyr172 (H172Y). Interactions of Si pocket(A) with N-terminus(B) are shown as solid lines and those of S1 pocket(B) with N-terminus(A) are shown as dashed lines. Similar disruption/destabilization was observed in the simulations of the nirmatrelvir-bound Mpro (Fig. SiO and Sii).

We next consider the H-bond between the imidazole of His172 and the N-terminal amine, which remained stable in the WT Mpro simulations (Fig. 2c and Fig. S3). An analogous H-bond for the H172Y mutant would be between the hydroxyl group of Tyr172 and the N-terminal amine. Indeed, this H-bond was occasionally sampled in all three runs at the beginning and it was completely abolished after 1 μs in run 1 and remained infrequently sampled in run 2 and rarely sampled in run 3 (Fig. 2c and Fig. S4-S6).

### A conserved aromatic stacking in the S1 pocket is destabilized in the simulations of free H172Y Mpro

The aromatic stacking between the absolutely conserved Phe140 and His163 is a key interaction that stabilizes the Mpro’s S1 pocket (Fig. 2a). This interaction was stable in the WT simulations, with the center-of-mass (COM) distance just below 4Å between the aromatic rings of Phe140 and His163 (Fig. 2e and Fig. S3). However, in simulation run 1 the aromatic stacking became lost after about 1 μs, with the stacking distance increased above 7Å (Fig. S4 and Fig. S7). The sudden breakage of the Phe140–His163 stacking was concurrent with a ~2-Å decrease in the COM distance between the oxyanion loop (residues 138–145)^4,18,19^ and Glu166 sidechain (Fig. S4) and a ~1-Å increase in the heavy-atom root-mean-square deviation (RMSD) of the oxyanion loop (Fig. S8). The latter is related to the decrease in the center of the mass distance between Glu166 and the oxyanion loop (Fig. S4), reminiscent of the oxyanion loop collapse observed in the simulations of the H172-protonated WT Mpro^15^ as well as an X-ray structure of SARS-CoV Mpro determined at pH 6 (PDB id 1uj1)^18^. In simulation run 2, the stacking interaction was stable until ~1.8 μs when the stacking distance increased by ~0.4 Å in protomer A; however in protomer B, the aromatic stacking was occasionally abolished, with the distance increasing beyond 15 A (Fig. S5). In simulation run 3, the Phe140–His163 stacking was stable in protomer A; however, it was abolished in protomer B for ~500 ns (stacking distance above 7 A) in the first 1 μs before the interaction was reestablished in the second 1 *μ*s; nonetheless, the COM distance occasionally increased to 9 Å (Fig. S6). Furthermore, in times when the stacking interaction was intact, the most probable COM distance between the two aromatic rings is increased by nearly 0.5 Å (Fig. 2e and Fig. S7). Thus, the simulations suggest that the H172Y mutation destabilizes the conserved Phe140–His163 interaction crucial for the stability of the S1 pocket.

### Destabilization of the aromatic stacking might be related to a nonnative H-bond between Phe140 and Tyr172

In comparing the H172Y and WT trajectories, we noticed that the hydroxyl group of Tyr172 can occasionally accept a H-bond from the backbone amide nitrogen of Phe140, whereas the analogous H-bond between the imidazole of His172 and the carbonyl of Phe140 is not possible. Interestingly, around the same time as the aromatic stacking between Phe140 and His163 became disrupted in the simulation run 1 of H172Y Mpro, the distance between the hydroxyl oxygen of Tyr172 and the amide nitrogen of Phe140 suddenly decreased (Fig. S4), which resulted in a significant increase of the H-bond occupancy from about 10% to about 45% (Fig. S9). A representative structure obtained from clustering analysis confirms a perturbed S1 pocket, whereby the Phe140–His163 stacking is abolished and Ser1* is moved away from the S1 pocket; however, Tyr172 is in a tight H-bond with the backbone of Phe140 (Fig. S10).

We hypothesized that a strong Phe140–Tyr172 H-bond would disrupt the aromatic stacking between Phe140 and His163. To test this hypothesis, we calculated the twodimensional probability densities of the Phe140–His163 and Phe140–Tyr172 distances. The density map shows a maximum located around the Phe140–His163 and Phe140– Tyr172 distances of 7.5 A and 3.0 Å, respectively (Fig. S10), representing a perturbed state in which the Phe140–His163 stacking is disrupted but a stable H-bond between Phe140– Tyr172 is formed. The density map also shows a local density maximum located at the Phe140–His163 and Phe140– Tyr172 distances of 4 Å and 3.1–3.6, respectively (Fig. S10), representing a state in which the aromatic stacking is intact and an occasional H-bond is formed between Tyr172 and Phe140. This analysis supports the hypothesis that the backbone interaction of Phe140 with Tyr172 destabilizes the sidechain interaction of Phe140 with His163, which may be responsible for the partial collapse of the oxyanion loop in run 1 (Fig. S4 and S8). However, since the complete disruption of the aromatic stacking was only observed in one of the three trajectories, this hypothesis requires further testing.

### Empirical energy calculations predicted decreased stability upon the H172Y mutation

Given the destabilization of the dimer interface and possibly the S1 pocket interactions, we wondered if the H172Y mutation destabilizes the Mpro. We addressed this question by calculating the folding free energy change upon mutation (ΔΔG_fold_) using the ddG_monomer application^20^ in the Rosetta software suite. Calculations (Fig. 3) for the Mpro dimers showed that the folding free energy of the mutant is about 9.9 ± 0.9 kcal/mol higher than the WT, mainly due to the destabilizing electrostatic (7.2 ± 1.0 kcal/mol) and H-bonding energies (4.8 ± 0.8 kcal/mol), and to a smaller extent the unfavorable van der Waals energies (2.2 ± 1.1 kcal/mol). Calculations for the Mpro monomers gave a similar ΔΔG_fold_ as for the dimer; the difference of 1.3 kcal/mol is within the error bar (Fig. 3). The contributions to the destabilization also come from the electrostatic, H-bonding, and van der Waals energies. This analysis suggests that the Mpro dimer destabilization upon mutation can be attributed to the destabilization of the monomers.

**Figure 3.**
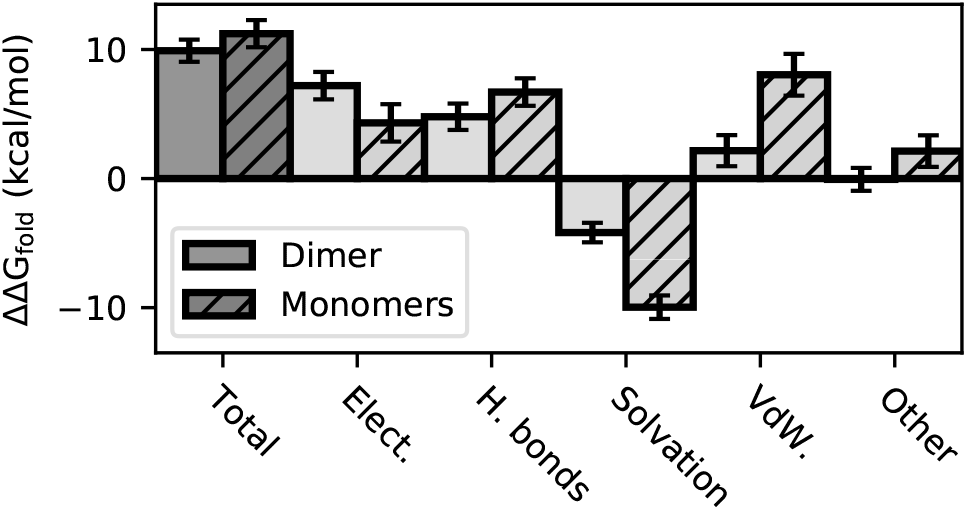
Changes in the folding free energy ΔΔG_fold_(WT → H172Y) calculated using the Rosetta ddG_monomer application^20^. The total ΔΔG_fold_ for the dimer (solid) and sum of monomers (striped) as well as the individual contributions are shown. Positive values indicate destabilization upon mutation.

Next we asked if the stability of the dimer interface is also affected by the mutation. If the mutation effect was restricted to the monomers, then the total ΔΔG_fold_ as well as the individual contributions would be the sum of the monomer energies. If however, ΔΔG_fold_ of the dimer is significantly higher (more positive) than that of the monomers, one could conclude that the dimer interface is destabilized. The similar total stability change for the dimer and sum of monomers does not suggest this is the case; however, the individual terms are different (Fig. 3). Most notably, the solvation energy of the dimer is less favorable than the monomers by 5.8 ± 2.0 kcal/mol, which balances out the less unfavorable van der Waals energy (5.9 ± 1.8 kcal/mol). Other terms are different as well, e.g., the electrostatic energy of the dimer is more unfavorable than the monomers (see later discussion). Thus, the energetics of the dimer interface is affected by the mutation although the net effect may be negligible.

To rationalize the above calculations, we examined the Rosetta generated structural models for the H172Y mutant and compared them with those for the WT Mpro dimer. The largest change is in Glu166, which upon losing the H-bond partner His172 is rotated away from Tyr172 (χ_3_ angle changed from −40° to −80 or 80° in the top three scored structures). This may explain the increased distance between Glu166 and the N-terminus of the opposite protomer (up to 0.5 A for the top three scored structures), which is consistent with the MD trajectories (Fig. 2c) and the more unfavorable electrostatic energy of the dimer as compared to the monomers upon mutation (Fig. 3). Replacing His172 with the larger Tyr172 also moved the Phe140 backbone amide nitrogen closer to the hydroxyl oxygen of Tyr172 (in comparison to the imidazole nitrogens of His172), with the distance of 3.35-3.65 Å between Phe140:N and Tyr172:OH in the top three scored structures. Although these distances do not indicate H-bonding, they do not exclude the possibility of transient (or strong) H-bond formation observed in the MD trajectories. As to the aromatic stacking between Phe140 and His162, the Rosetta generated structures showed an increase of 0.15 A between the COM of the two rings in the mutant, which, albeit small, is consistent with the destabilization observed in the MD trajectories.

### S1 pocket interactions with N-terminus* are also destabilized in the simulations of the nirmatrelvir bound Mpro

To probe the effect of H172Y mutation on the Mpro’s affinity for nirmatrelvir, we first performed 2-μs simulations of the WT and H172Y Mpros in complex with nirmatrelvir (Table S1). In these simulations, nirmatrelvir remained stably bound with the Mpro (Fig. S1) and the aromatic stacking between Phe140 and His163 was intact; however, similar to the free Mpro, the N-terminus interaction with Phe140 was completely lost and those with Glu166 and Tyr172 were significantly weakened in both protomers (Fig. S11 and S12). Surprisingly, the RMSD of the oxyanion loop was unstable (Fig. S8). These data are consistent with the simulations of the free H172Y Mpro, and suggest that the S1 pocket in the inhibitor-bound form is also destabilized by the mutation.

### Perturbation of the P1 site and formation of the Phe140–Tyr172 nonnative hydrogen bond in the simulations of the nirmatrelvir-bound Mpro

To further probe the stability of nirmatrelvir binding, we compared the distributions of nirmatrelvir’s RMSD with respect to the X-ray structure (PDB id 7vh8)^5^ calculated from the trajectories of the WT and H172Y Mpros. The peak is slightly right shifted for the H172Y relative to the WT simulations (Fig. S13), which indicates that nirmatrelvir has a small conformational change when complexed with the mutant Mpro. Calculations of the atom-based protein-ligand contact distances showed that the change mainly affects the γ-lactam ring in the P1 position, whereby the amide nitrogen forms a H-bond with the carboxylate oxygen of Glu166 in the X-ray structure (PDB id 7vh8)^5^. This H-bond was stable in the WT simulations, with an occupancy over 60%, but it was significantly weakened in the H172Y simulations, with an occupancy about 20% (Fig. S13). On the other hand, the H-bond between the lactam nitrogen and the backbone carbonyl oxygen of Phe140 was stabilized in the mutant simulations, with an occupancy increase of about 30% as compared to the WT simulations (Fig. S13). This analysis is consistent with the representative structure from the clustering analysis of the H172Y simulations, which showed that the H-bond between the lactam nitrogen and Glu166 is absent (Fig. S13). Importantly, similar to the ligand-free simulation run 1, the nonnative H-bond between Tyr172 and Phe140 is formed (Fig. S13). This consistency suggests that the perturbation of the S1 pocket by the H172Y mutation is responsible for the change in the P1 site binding, which we speculate may contribute to the decreased affinity for nirmatrelvir.

### Free energy simulations and empirical calculations predict decreased nirmatrelvir affinity for the H172Y Mpro

To further examine the mutation effect on the affinity of nirmatrelvir-Mpro noncovalent binding, we calculated the binding free energy change upon mutation, which according to the thermodynamic cycle is the same as the difference in the mutation free energies of the free and ligand-bound forms (Fig. 4, top). We applied two methods to calculate the mutation free energies. First, we conducted the alchemical free energy perturbation (FEP)^21^ simulations using the implementation^22,23^ in the NAMD2 package^24^. Both the WT-to-mutant and mutant-to-WT transformations were performed, although the latter may be less accurate due to the use of the computationally mutated structure. Both transformation predicted the mutant to have a significantly decreased binding affinity (Fig. 4, bottom). The more reliable WT-to-mutant transformation gave the values of 2.7±0.19 kcal/mol for protomer A and 2.2±0.17 kcal/mol for protomer B. We also applied an empirical approach to calculate the mutation free energies using Rosetta’s flex ddG protocol^25^. These calculations also predicted a lower binding affinity for the mutant, although to a smaller extent (about 0.3 kcal/mol) as compared to the more accurate FEP calculations.

**Figure 4.**
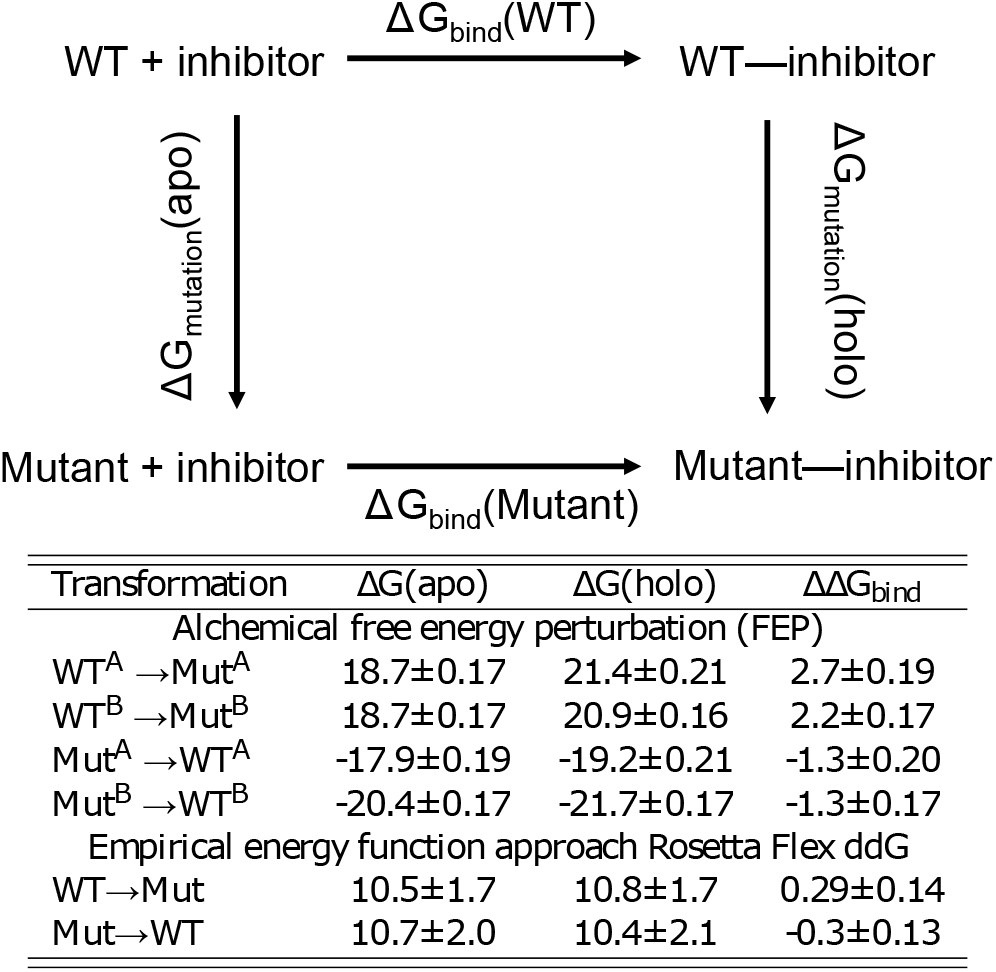
Top. Thermodynamic cycle used to calculate the difference in the noncovalent binding free energy between the mutant and WT Mpros: ΔG_bind_(Mutant) - ΔG_bind_(WT) = ΔG_mutation_(holo) - ΔG_mutation_(apo). Bottom. FEP and empirical calculations of the change in nirmatrelvir binding free energy of Mpro upon the H172Y mutation. For FEP calculations, transformations were performed on each protomer separately, whereas Rosetta calculations transformed H172 in both protomers simultaneously. ΔΔG_bind_ calculated from the transformation from mutant to WT is less accurate, as it was initiated from the modeled mutant structure. Each calculation was repeated a number of times (see Table S2), the mean and standard errors are reported.

### Artificial neutral network identified conformational changes of the oxyanion loop region upon mutation

To further analyze the MD trajectories to discern the mutation effect on the conformational dynamics of the Mpro, we utilized a newly developed artificial neural network called DiffNets^26^, which makes use of autoencoder and classifier to detect structural differences of protein variants based on MD trajectories (Fig. 5a). We created two DiffNets to discern the effect of H172Y mutation on the free and ligand-bound Mpros. Positions of C, CA, CB, and N atoms of the WT and H172Y Mpros were fed as input into two encoders, with atoms near the mutation site fed into a separate encoder. These positions were then encoded to a latent (reduced dimensional) space followed by reconstruction to reproduce the input positions (Fig. 5a and Fig. S14). A classifier was applied to the latent space to determine if the frame comes from a WT or H172Y trajectory (Fig. 5b). clustering was then applied in the latent space to identify pairwise distances that are most correlated (largest R^2^ values) with the predicted labels (WT vs. H172Y). Interestingly, for both the free and ligand-bound forms, the Cα distances most correlated with the labels involve Gly138, which is the first residue of the oxyanion loop (residues 138–145, Fig. 5c and Table S4). The distance from Gly138 to Ser144 is 1 Å greater in the H172Y relative to WT trajectories of the ligand-bound form (Fig. 5d). This distance change is consistent for both protomers in both trajectories (Fig. S15). In the free Mpro, the shift in the Gly138-Ser144 distance based on the aggregated trajectories and protomers is subtle (Fig 5d); however, the shift is very pronounced for protomer B in two (out of three) trajectories (Fig. S15). The distance from Gly138 to Thr135, which is in the unstructured region preceding the oxyanion loop, is also greater by about 1 Å for the H172Y vs. WT trajectories in both free and ligand-bound enzyme forms (Fig. 5d and Fig. S16). These data suggest that the oxyanion loop region is more extended upon the H172Y mutation.

**Figure 5.**
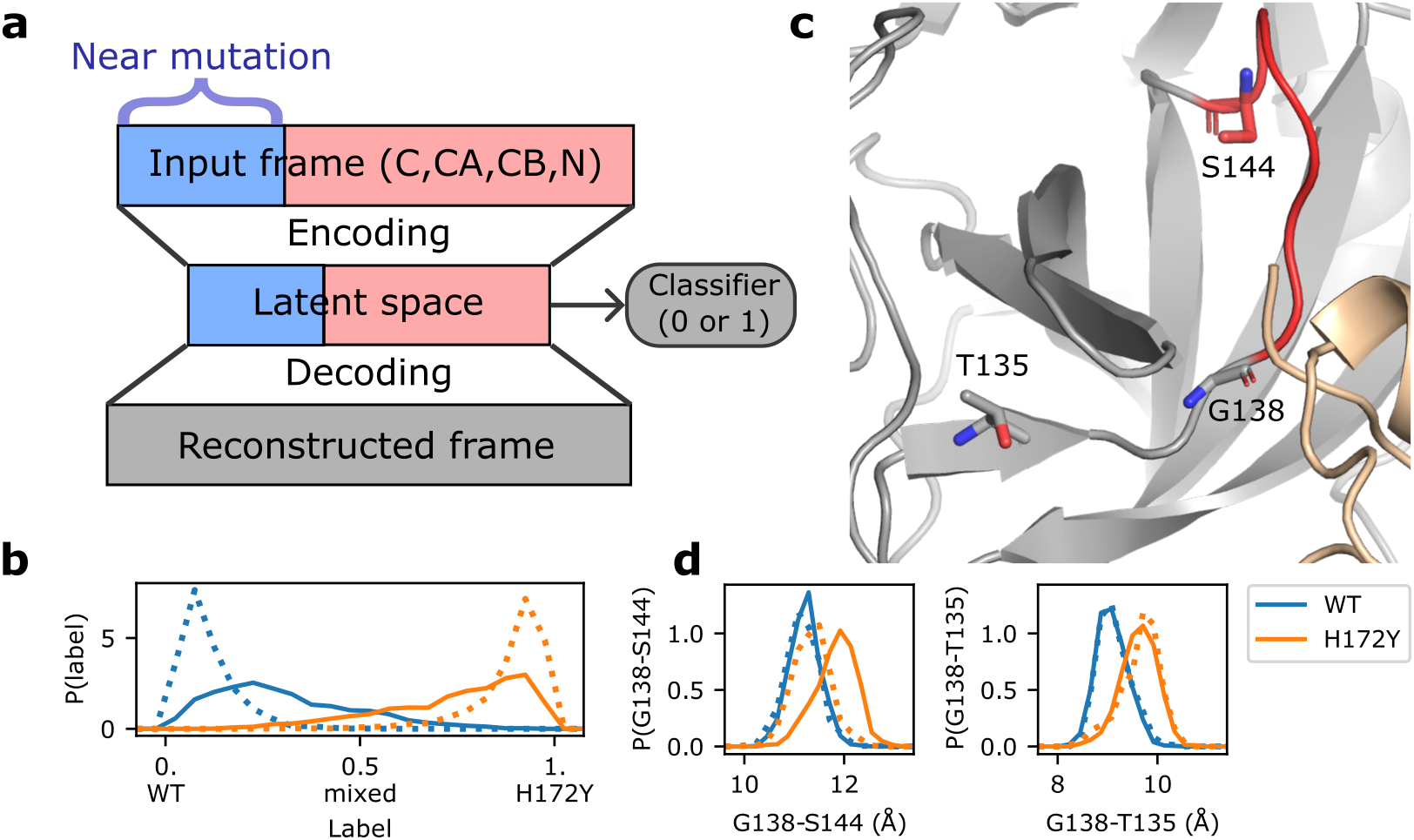
Artificial neural network detects mutation-induced conformational changes to the oxyanion loop. a. Schematic architecture of the autoencoder DiffNets^26^. which was used to detect differences between protein structures from two trajectories. b. Classification (WT vs. H172Y) of the free (dotted) and ligand-bound (solid) trajectories. The three free and two ligand-bound WT (blue) or H172Y (brown) trajectories were aggregated. c. Zoomed-in view of the oxyanion loop (red, residues 138—145; among them 143—145 form the oxyanion hole) and the three residues (licorice) involved in the important distances (Gly138–Ser144 and Gly138–Thr135) that distinguish between the WT and H172Y trajectories (i.e., highly correlated with the predicted labels). The oxyanion hole is comprised of Gly143, Ser144, and Cys145. d. Probability distributions of the Cα distances from Gly138 to Ser144 (left) and Thr135 (right) from the free (dotted) or ligand-bound (solid) WT (blue) and H172Y (brown) Mpro trajectories. The aggregate trajectories including both protomers were used. Data for individual trajectories and protomers as well as other distances involving Gly138 are given in Figures S14 and S15.

### Experiments confirm that the H172Y mutation reduces Mpro’s stability, catalytic activity, and susceptibility to nirmatrelvir

Following the simulation study, we measured the thermal stability and enzyme kinetics of WT and H172Y Mpros as well as the IC_50_ values of nirmatrelvir (Fig. 6). Thermal-shift assays were used to determine the unfolding temperatures (*T_m_*) of the WT and mutant Mpros (Fig. 6a). The *T_m_* for the WT was found to be 58.11 °C, whereas that of the H172Y Mpro was lower by 4.16°C (Fig. 6d), indicating a destabilization of the enzyme. According to an empirical formula^27^ 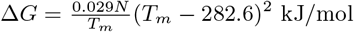, where N is the number of residues and *T_m_* is in Kelvin, the decrease of T_m_ corresponds to roughly 2.3 kcal/mol decrease of unfolding free energy, which is in qualitative agreement with the prediction by the Rosetta calculation (Fig. 3).

**Figure 6.**
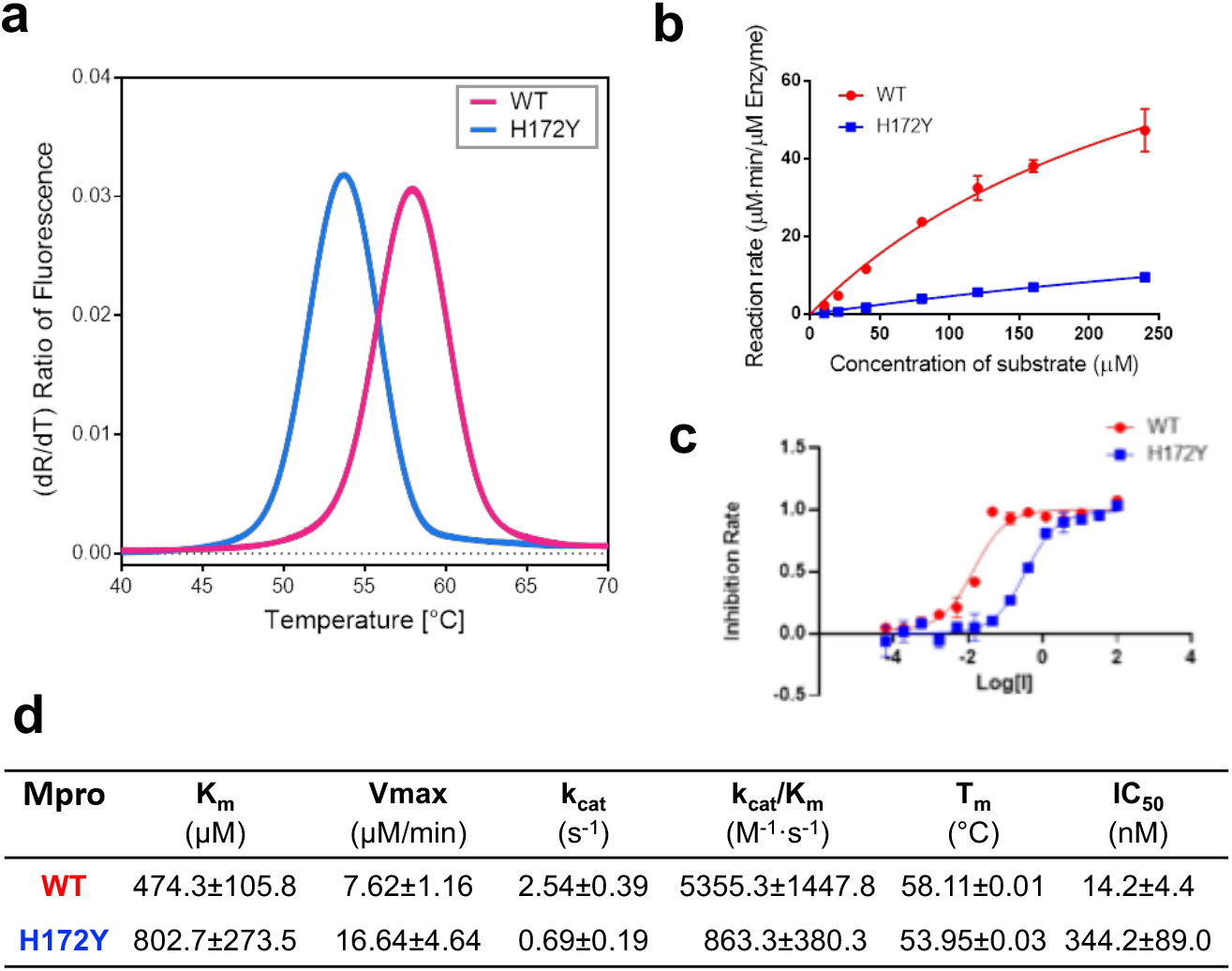
H172Y Mpro has reduced thermal stability, enzyme activity, and susceptibility to nirmatrelvir as compared to the WT. **a.** Melting curves of the WT (red) and H172Y (blue) Mpros based on the temperature profile of the first derivative of the ratio of the autofluorescence at 350 and 330 nm. **b.** Reaction rate vs. substrate concentration for the WT (red) and H172Y (blue) Mpros using the FRET assay. **c.** Inhibition rate of nirmatrelvir vs. its concentration (μM) for the WT (red) and H172Y (blue) Mpros. **d.** Summary of the kinetic constants, melting temperatures of the Mpros, and the IC_50_ values of nirmatrelvir.

The reaction rate measurement using the FRET assay revealed a significant decline in the catalytic efficiency for the H172Y relative to the WT Mpro (Fig. 6b and d). The *k*_cat_/*K*_m_ value obtained for the WT enzyme is 5355.3 M^-1^s^-1^, while that for H172Y is 863.3 M^-1^s^-1^, i.e. only 16% enzyme activity remains in the FRET assay, compared to the WT. The *k_cat_* value (enzyme turnover number) for H172Y is 0.69 s^-1^, which is only 27% of the WT value. The Michaelis constant *K*_m_ value obtained for H172Y is 802.7 μM, which is 69% larger than the value for the WT, indicating that the mutation significantly reduces the affinity for substrate binding.

The significant decrease in the enzyme efficiency (decreased turnover number and substrate binding) may be explained by the extension of the oxyanion loop (increase of the distance between Gly138 and Ser144) (Fig. 5d). The latter may be attributed to the loss of the N-terminus interaction with Phe140 (Fig. 2b). Since the oxyanion loop forms the wall of the S1 pocket, a subtle conformational change may reduce the substrate binding affinity. Since the oxyanion hole residues Gly143 and Cys145 directly interact and stabilize the reaction intermediate^18^, this conformational change may also perturb the transition state and consequently the kinetics of enzyme catalysis.

The inhibitory activity of nirmatrelvir was followed using the FRET assay (Fig. 6c and d). The IC_50_ against the H172Y mutant is 344.2±89.0 nM, which is 24.2 times higher than that for the WT protein. Converting the IC_50_ values to *K_i_* values using the web server^28^ gave a similar ratio of 24.5 times for the *K_i_* values of H172Y vs. WT, which corresponds to a free energy change of about 1.9 kcal/mol. This reduction of binding affinity by 1.9 kcal/mol is in good agreement with the FEP estimated values of 2.3–2.7 kcal/mol and consistent with the empirical calculations although the magnitude of ΔΔG_bind_ is much smaller (about 0.3 kcal/mol, Fig. 4 bottom).

### Additional simulations based on a X-ray structure model of H172Y Mpro

In the final stage of the manuscript preparation, we performed an additional fixed-charge MD simulation based on an unpublished preliminary X-ray structure model of H172Y Mpro (Table S2). During the 2-μs simulation, the N-terminus h-bond/salt bridge interactions with Phe140 and Glu166 were significantly destabilized in protomer A and completely disrupted in protomer B (Fig. S16). In protomer B, the disruption of the S1 pocket-N-terminus interactions is concomitant with a small (about 0.3 Å) increase in the Phe140–His163 aromatic stacking distance (Fig. S16) and a nearly 1-Å increase in the RMSD of the oxyanion loop (Fig. S17). Consistently, the distances between Gly138 and Ser144/Thr135 in both protomers are increased (by about 1 Å) compared to the WT, consistent with the simulations based on the computationally mutated H172Y Mpro structure (Fig. S18). In fact, for protomer B, the distributions of the two distances are very similar between this new simulation and those based on the modeled structure of the H172Y Mpro. Thus, the additional simulation based on a different starting structure confirmed the disruption of the S1 pocket-N-terminus^*^ interactions, the conformational change of the oxyanion loop, and the destabilization of the Phe140–His163 stacking.

### Our MD data are consistent with the new X-ray structures of H172Y Mpro

As we were preparing the manuscript, a bioRxiv paper by Hu et al.^10^ was published that reports the X-ray structures of the free and inhibitor GC-376-bound H172Y Mpros. In the ligand-free X-ray structure (PDB id: 8d4j)^10^, the salt bridge between Glu166 and the N-terminus^*^ is lost in one protomer, and the H-bond between Phe140 and the N-terminus^*^ is lost in both protomers (Table S3). These data corroborate the simulation finding of the abolished interactions between Phe140/Glu166 and the N-terminus^*^ (Fig. 2). Note, in the GC376-bound structure (PDB id: 8d4k)^10^, the position of Ser1 is not resolved. Another agreement between simulation and the reported X-ray structures of H172Y Mpro is with regards to the increased Ca distances of Gly138-Ser144 and Gly138-Thr135. They are respectively 0.2/0.2 and 0.2/0.5 Å greater in the free/GC376-bound H172Y (PDB 8d4j/8d4k)^10^ as compared to the WT Mpro (PDB 7vh8)^5^ structure. Thus, the X-ray structure models are in support of a mutation-induced conformational change of the oxyanion loop.

### The present biochemical data are consistent with the reported data

Hu et. al^10^ reported that the *k*_cat_/*K*_M_ value of the H172Y Mpro is 13.9-fold lower than the WT, compared to the 6.2-fold decrease determined in this work, thus confirming a significant reduction in the catalytic efficiency upon mutation. We note that the impact of mutation on Mpro’s cleavage activity likely varies depending on the substrate. Thus, the difference in the *k*_cat_/*K*_M_ value reduction may be due to the different FRET substrate used in the experiments. Hu et. al^10^ also reported that the Ki value of nirmatrelvir is >113.7 fold higher for the H172Y than the WT Mpro, compared to the roughly 24.5-fold increase in the *K*_i_ value estimated^28^ from the 24-fold increase in the IC_50_ value determined in this work. Thus, both experiments confirmed a significantly reduced affinity and activity of nirmatrelvir against H172Y relative to the WT Mpro.

## Concluding Discussion

Employing an in silico structure model and a battery of state-of-the-art computational techniques, including constant pH and fixed-charged MD, alchemical free energy simulations, empirical energy calculations as well as artificial neural networks, we made prospective predictions regarding how structure, dynamics, folding stability, and inhibitor binding of SARS-CoV-2 Mpro change upon the H172Y mutation. The MD simulations of the free and nirmatrelvir-bound Mpros showed that the mutation disrupts or significantly destabilizes the interactions between the S1 pocket residues Phe140, Glu166, and His172 and the N-terminus of the opposite protomer. The conserved aromatic stacking between Phe140 and His163 in the S1 pocket was also destabilized upon mutation. The analysis using artificial neural network found that the oxyanion loop is extended for both free and ligand-bound H172Y Mpros. Remarkably, these results are in agreement or consistent with the newly reported X-ray structures of the free and GC376-bound H172Y Mpro (PDB ids: 8d4j and 8d4k)^10^ as well as our preliminary structure model of the H172Y Mpro (Hilgenfeld and coworkers, unpublished). The simulation data may explain the significant (84%) reduction in the *k*_cat_/*K_m_* value due to the H172Y mutation. In particular, the conformational change of the oxyanion loop that stabilizes the reaction intermediate may explain the significant (73%) decrease in the *k*_cat_ value (decreased enzyme turnover number), although quantum mechanical/molecular mechanics (QM/MM) calculations may offer more detailed clue regarding the perturbation of kinetics. The destabilization of the S1 substrate pocket as well as the change of the oxyanion loop may explain the significant (69%) increase in the *K_m_* value which represents the decreased substrate binding affinity.

The Rosetta^20^ predicted folding stability decrease upon mutation is consistent with the reduced *T*_m_ value determined using the thermal-shift assays. The energy analysis suggested that the stability decrease is largely due to the unfavorable change of the electrostatic and H-bond energies of the monomers, consistent with the MD data. Both the Rosetta energy calculations^25^ and the more accurate FEP simulations predicted that the H172Y Mpro has a reduced binding affinity for nirmatrelvir, which is consistent with the significant increase in the IC_50_ value determined by us or the K_*i*_ value determined by Wang and coworkers^10^ and by Pfizer’s disclosure^1^. The simulation data suggested that the decreased binding affinity between nirmatrelvir and the H172Y Mpro may be attributed to the dynamical perturbation of the S1 pocket, which weakens the H-bond between Glu166 and the γ-lactam nitrogen in the P1 position. The MD data suggests that Phe140 plays a critical role here, as its interaction with the N-terminus^*^ is completely abolished, which may drive the conformational change of the oxyanion loop. The perturbation to Phe140 may also explain the weakened stacking interaction with His163 and the nonnative H-bond formation with Tyr172. The destabilization of the interaction between Glu166 and the N-terminus^*^ may be a major contributor to the weakened interaction with the γ-lactam nitrogen of nirmatrelvir at the P1 position. This finding also suggests that optimization of the γ-lactam moiety may offer a route to improve the antiviral potency. rescue function while maintaining antiviral resistance (e.g., as demonstrated by a recent experiment^10^). The recent explosion of COVID-19 cases in China and wide-spread use of nirmatrelvir therapy in the US raise the odds of resistance mutations. On that note, our work demonstrates the predictive power of combined molecular simulation and artificial intelligence approaches, and together with biochemical experiments they provide an important tool for the active surveillance of continually emerging SARS-CoV-2 Mpro mutations and the discovery of new antiviral inhibitors. The presented workflow is general and can be applied to characterize mutation effects on any protein drug targets.

### Computational methods and protocols

#### Structure preparation for molecular simulations

The Modeller software^11^ was used to generate an initial structural model of the H172Y mutant of SARS-CoV-2 Mpro, with the X-ray crystal structure of the wild-type (WT) Mpro in complex with nirmatrelvir (PDB id 7vh8)^5^ as a template. Next, the WT and H172Y Mpro were prepared for MD simulations using the LEAP utility of Amber^17^, with the termini left free. The protein was represented by the Amber ff14SB force field^29^ and water molecules by the TIP3P model^30^.

#### Continuous constant pH molecular dynamics (CpHMD) titration simulations

The protonation states of the mutant H172Y Mpro were determined using the GPU-accelerated GBNeck2 implicit solvent CpHMD titration^13^ with asynchronous pH replica exchange^14^, as in the previous CpHMD simulations of the WT Mpro^15^. Briefly, 9 replicas were used over pH range 5 to 9 with an interval of 0.5 pH unit. Each replica was simulated at 300 K with an ionic strength of 0.15 M and an effectively infinite cutoff (999 Å) for nonbonded interactions. Each replica was run for 30 ns, with an aggregate sampling time of 270 ns. All sidechains of Asp, Glu, His, Cys, and Lys were allowed to titrate, with the model *pK*_a_’s of 3.8, 4.2, 6.5, 8.5, and 10.4, respectively. More details of the simulation settings are given in Ref.^15^

#### Fixed-charge molecular dynamics simulations

The octahedral water box was used to solvate the protein, with a distance of at least 11 Å between the protein heavy atoms and the water oxygen atoms at the edges of the box. Sodium and chloride ions were added to neutralize the system and create an ionic strength of 150 mM. For the nirmatrelvir-bound Mpro simulations, the reversible bound model in the X-ray structure was used. The ligand parameters were generated using the general Amber force field (GAFF2) with partial charged derived using the AM1 BCC method ^31,32^ All simulations were carried out using Amber20^17^. First, energy minimization with a harmonic restraint of 100 kcal/mol/Å^2^ on the protein heavy atoms was performed for 10000 steps using the steepest descent algorithm followed by 10000 steps using the conjugate gradient algorithm. Next, the system was heated from 100 K to 300 K using the same harmonic restraint in the canonical ensemble by 1 ns. Five equilibration stages using harmonic forces of 10, 5, 2, 1, and 0.1 kcal/mol/Å^2^ were then performed for 50 ns in the NPT ensemble. The pressure was maintained at 1 atm using the Berendsen barostat with a relaxation time of 0.1 ps, and the temperature was maintained at 300 K using the Langevin thermostat with a collision frequency of 1.0 ps^-1^.^17^ The particle-mesh Ewald^33^ method was used to treat the long-range electrostatics with a grid spacing of 1 Å. A cutoff of 8 Å was used for van der Waals interactions. SHAKE was used to increase the time step to 2 fs. Finally, the production simulations were performed for 2 μs for both the ligand-free and nirmatrelvir-bound WT and H172Y Mpros. A summary of the simulations is given in Table S2.

#### Trajectory analysis using artificial neural network

We applied DiffNets^26^, an artificial neural network with a split-autoencoder architecture for detecting structural differences between the MD trajectories of protein variants. DiffNets^26^ follows the autoencoder architecture: an encoder collapses the high dimensional input into a (low dimensional) latent space, then a decoder reconstructs the points in latent space back to the original input (Fig. 5). Two additional functions are added. First, the user labels trajectories either 0 or 1, based on a binary quantity (activity, mutation, etc.); the latent space is then used by a classifier in order to predict the input label. This classifier is trained alongside the encoding and decoding layers and the predicted label is utilized in the loss function in order to separate the labels on the latent space. Second, the atomic coordinates are separated based on their proximity to the mutation site and fed into separate encoding layers. The resulting latent variables are then concatenated to form the full latent space used by the decoder.

Two separate DiffNets were built on either the free or the ligand-bound WT and H172Y fixed-charge trajectories (Table S2). First, the trajectories of WT (two) and H172Y (three) were strided every 1 ns to generate frames for each protein (2,000 ×3 for the free, 2,000×2 for the ligand-bound state) followed by the extraction and alignment of the coordinates of the N, C, CA, and CB atoms (2398 total). After the mean was subtracted from each frame and the trajectory was whitened, the frames were used to train a split-autoencoder, where the atoms within 10 Å of any atom of H/Y172 (approx. 700 atoms) were fed to a supervised encoder while the rest of the protein was fed to a second unsupervised encoder. For the supervised autoencoder, a classification task (label 0 for WT and 1 for H172Y mutation) was added to the latent space. 90% of frames were used for training, while 10% were reserved for testing. Both encoders encode the positions of the input frame into a latent space that is joined to from a vector of 50 components. This latent space is then clustered into 200 clusters using a k-centers/k-mediods hybrid algorithm, and the centroid of each cluster is decoded to produce a reconstructed representative frame. These frames are then used to calculate pairwise distances between CA atoms within 15 Å of either mutation site. The correlation between each distance and the predicted label of the frame was calculated, and the top 10 most correlated distances (with *R*^2^ values ranging 0.93-0.85) were designated significant distances (Table S3) and visualized using PyMOL^34^. The significance of these distances was verified by plotting and comparing the distributions of the real distances from the trajectory frames of the WT and H172Y Mpros. Further details of the protocol are given in Ref ^26^. The expectation maximization algorithm which adjusts target labels during training was turned off, as it is not relevant for our task of interest which is to use latent space to recognize structural features related to the classification labels (WT vs. mutant).

#### Empirical calculations of protein stability changes

The changes of the folding free energies (ΔG_fold_ = -ΔG_stability_) of the apo Mpro dimer and monomers upon mutation was calculated using the ddg_monomer application^20^ within the Rosetta software suite. In this method, an ensemble of structure models of the mutant was generated from the input WT structure (PDB id 7vh8, with nirmatrelvir removed)^5^. The change in the folding free energy due to mutation (ΔΔG_fold_) was calculated as the difference in the Rosetta energies between the WT and mutant structures. A positive value indicates a decreased stability from the mutation. The high-resolution protocol (with both backbone and sidechain relaxation) was followed^20^. First, Rosetta’s standard side-chain optimization module was used to optimize the input WT structure (PDB 7vh8^5^); then three sequential minimization calculations were performed where the Lenard-Jones potential was scaled by 0.1, 0.33, and 1.0 respectively. Distance restraints on C_alpha_ atoms were applied to prevent the backbone from deviating from the initial structure. This process was repeated 50 times for both the WT and (generated) H172Y Mpro dimer structures, then the average score for each system was calculated using the REF2015 energy function^35^. This calculation was also performed using monomeric Mpro, where the second chain of 7vh8 was removed.

#### Calculation of ligand binding free energy changes using free energy perturbation (FEP)

The alchemical FEP method^21,36^ was used to calculate change in the noncovalent binding free energy going from the WT to the H172Y mutant Mpro: ΔΔG_bind_ = ΔG_bind_ (Mutant) —ΔG_bind_(WT), which according to the thermodynamic cycle (Fig. 4) can be calculated from the difference in the mutation free energies: ΔΔG_bind_ = ΔG_mutation_(holo) — ΔG_mutation_(apo). The last two terms can be calculated via FEP as the free energies of transforming His172 to Tyr172 in the ligand-bound and ligand-free forms. Note, the binding free energy difference is related to the ratio of the *K_d_* values, ΔΔG_bind_ = —RTln(K_d,Mut_/K_d_,W_T_).

The FEP simulations were performed using NAMD2^22–21^. The X-ray structure of WT Mpro in a noncovalent complex with nirmatrelvir (PDB id 7vh8, the noncovalent binding mode)^5^ was used to create a model H172Y mutant as in the fixed-charge simulations. The proteins were represented by the CHARMM36m force field^37,38^, and the noncovalently bound nirmatrelvir was represented by the CGenFF force field obtained through the Paramchem server^39,40^. To allow an integration timestep of 2 fs, all bonds and angles involving hydrogen atoms were constrained using the SHAKE algorithm^41^. The temperature was maintained at 310 K by Langevin dynamics with a damping coefficient *γ* of 1 ps^-1^, and the pressure was controlled at 1 atm by the Nosé-Hoover Langevin piston method^42,43^. The van der Waals interactions were smoothly switched off from 10 to 12 Å using a switching function. The particle mesh Ewald (PME) method^33^ was used to calculate long-range electrostatic energies with a sixth-order interpolation and a grid spacing of 1 Å. Each starting structure (WT or mutant Mpro in complex with nirmatrelvir) was equilibrated for a total of 52 ns whereby the protein and ligand were restrained in the initial 2 ns simulation (0.25 ns with heavy atom restraint at 2.5 kcal/mol Å^2^, 0.25 ns with heavy atoms of the protein and ligand restrained at 1.25 kcal/mol Å^2^, 0.5 ns with backbone atoms of protein and heavy atoms of the ligand restrained at 1.25 kcal/mol Å^2^, and 1 ns with *C_α_* atoms of the protein and heavy atoms of the ligand restrained at 1.25 kcal/mol Å^2^). The equilibrated structure was used for the FEP simulations. The starting structures for the apo FEP simulations were generated by deleting the ligand from the 50-ns equilibrated holo structures. In total, there were 8 simulation sets: 2 holo wild type, 2 holo mutant, 2 apo wild type, and 2 apo mutant. The hybrid H/Y172 complexes, in which the mutated residue 172 comprising the imidazole and phenol rings representing as the appearing or disappearing particles, were modeled using VMD ^44^.

The progress of the alchemical transformation was described by the coupling parameter λ, which was gradually scaled from 0 to 1 for the forward (e.g., His to Tyr or Tyr to His) and from 1 to 0 for the backward transformation (e.g., Tyr to His or His to Tyr). In each simulation set, the backward transformation was performed consecutively from the forward transformation. A transformation simulation lasted 12 ns, comprising 20 intermediate λ states/windows. The sampling of each window lasted 0.6 ns, with the last 0.5 ns used for ensemble averaging. The aggregate simulation time was 192 ns. The electrostatics interactions of the disappearing particles were linearly decoupled from the system from λ = 0 to λ = 0.5, while those of the appearing particles were linearly coupled from λ = 0.5 to λ = 1. For the van der Waals interactions, a soft-core potential was also applied to ensure a gradual transformation. The disappearing particles were fully coupled at λ = 0 and fully decoupled at λ = 1, while the appearing particles were fully decoupled at λ = 0 and fully coupled at λ = 1. The ParseFEP toolkit^45^, implemented in VMD was used to test convergence and calculate the transformation free energies. The latter was estimated using the Bennett acceptance ratio (BAR) method^46,47^.

#### Empirical calculation of ligand binding free energy changes with Rosetta

The change in nirmatrelvir binding free energy due to the H172Y mutation was also studied using the flex ddG protocol^25^ in the Rosetta modeling software (version 2017.52.59948). Although designed for the prediction of changes in protein-protein binding affinities upon mutations, a recent benchmark study^48^ found that the flex ddG protocol is able to quantitatively predict changes in protein-ligand binding affinities upon mutations. The flex ddG protocol calculates the binding energies using the Rosetta energy function^35^ and the “backrub” protocol ^49^, which performs Monte-Carlo trials to sample local sidechain and backbone conformational changes near the mutation site. The calculations for the forward mutations, ΔG_WT→Mut_(apo) and ΔG_WT→Mut_(holo), were based on the X-ray structure of the WT Mpro dimer (PDB id 7vh8)^5^, while the calculations for the backward mutations, ΔΔG_Mut→WT_(apo) and ΔΔG_Mut→WT_(holo), were based on the computationally mutated structures (using Modeller^11^ and the PDB 7vh8^5^). Parameters for nirmatrelvir were obtained using the molfile_to_params.py script in Rosetta^25^. The protocol was repeated 40 times, with 35,000 backrub trials for each run. The final trial of each run was scored using the Rosetta Energy function 2015 (REF201 5) ^35^. The average energy score for each model (WT or mutant, apo or holo) was calculated and the change in binding free energy upon mutation was estimated using the thermodynamic cycle (Fig. 4) as ΔΔG_bind_ = [E_Mut_(holo) — E_WT_(holo)] — [E_Mut_ (apo) – E_WT_(apo)], where *E* represents the energy score.

### Protein production and characterization

#### Cloning of SARS-CoV-2 Mpro H172Y

The H172Y mutation was inserted by overlap extension-PCR reaction. A pair of special primers, H172Y_forward (ACTGGTGTA<u>TAT</u>GCCGGGACGGACT; the underlined sequence corresponds to the mutated H172Y codon) and H172Y.reverse (AGTCCGTCCCGGCAT<u>ATA</u>CACCAGT) were designed. The first PCR reaction was performed to generate two splice fragments containing a 5’ overhang. The WT Mpro coding gene with BamHI and XhoI sites was amplified from the Mpro construct as described previously^50^, and was used as template. The second PCR joined these two spliced fragments to generate the PCR product encoding the H172Y mutated Mpro including the cleavage sites of the restriction enzymes for cloning into the vector PGEX-6p-1 (GE Healthcare). The amplified PCR product was digested with BamHI and XhoI and ligated into the vector PGEX-6p-1 digested with the same restriction enzymes. The gene sequence of the Mpro was verified by sequencing (MWG Eurofins).

The sequence-verified SARS-CoV-2 Mpro construct was transformed into E. coli strain BL2 (DE3) (Novagen). Transformed clones were pre-cultured at 37° C in 50 mL 1 x YT medium with ampicillin (100 μg/mL) for 3 h, and the incubated culture was inoculated into 4 L 1 x YT medium supplied with 100 μg/mL ampicillin. 0.5 mM isopropyl-D-thiogalactoside (IPTG) was added for induction of the overexpression of the Mpro gene at 37°C when the OD600 reached 0.8. After 5 h, cells were harvested by centrifugation at 9954 x g, 4°C, for 15 min. The pellets were resuspended in 30 mL buffer A (20 mM Tris, 150 mM NaCl, pH 7.8; pH of all buffers was adjusted at room temperature) and then lysed by sonication on ice. The lysate was clarified by ultracentrifugation at 146,682 x g at 4°C for 1 h. The supernatant was loaded onto a HisTrap FF column (GE Healthcare) equilibrated with buffer A. The HisTrap FF column was washed with 150 mL buffer A to remove unspecifically bound proteins, followed by elution using buffer B (20 mM Tris, 150 mM NaCl, 500 mM imidazole, pH 7.8) with a linear gradient of imidazole ranging from 0 mM to 500 mM, 20 column volumes. The fractions containing target protein were pooled and mixed with PreScission protease at a molar ratio of 5:1 and dialyzed into buffer C (20 mM Tris, 150 mM NaCl, 1 mM DTT, pH 7.8) at 4°C overnight, resulting in the target protein with authentic N- and C-termini. The PreScission-treated Mpro was applied to connected GST-trap FF (GE Healthcare) and nickel columns to remove the GST-tagged PreScission protease, the His-tag, and protein with uncleaved His-tag. The His-tag-free Mpro in the flowthrough was concentrated by using Amicon Ultra 15 centrifugal filters (10 kD, Merck Millipore) at 2773 x g, and 4°C. The protein was loaded onto a HiLoadTM 16/600 SuperdexTM 200pg column (GE Healthcare) equilibrated with buffer A. Fractions eluted from the Superdex200 column containing the target protein with high purity were pooled and subjected to buffer exchange (20 mM Tris, 150 mM NaCl, 1 mM EDTA, 1 mM DTT, pH 7.4).

#### Determination of protein stability of SARS-CoV-2 Mpro WT and H172Y by nano differential scanning fluorimetry (nanoDSF)

Thermal-shift assays of SARS-CoV-2 Mpro and its H172Y mutant were carried out using the nanoDSF method as implemented in the Prometheus NT.48 (NanoTemper Technologies). The nanoDSF method is based on the autofluorescence of tryptophan (and tyrosine) residues to monitor protein unfolding. As the temperature increases, the protein will unfold and the hydrophobic residues of the protein get exposed, the ratio of autofluorescence at wavelengths 350 nm and 330 nm will change. The first derivative of 350/330 nm can be used to determine the melting temperature (*T_rmm_*). 30 μM of WT or mutant protein were diluted in a final volume of 15 μL reaction buffer containing 20 mM HEPES, 120 mM NaCl, 0.4 mM EDTA, 4 mM DTT, 20% glycerol, pH 7.0. Then the proteins were loaded onto Prometheus NT.48 nanoDSF Grade Standard Capillaries (PR-C002, NanoTemper Technologies), the fluorescence signal was recorded under a temperature gradient ranging from 25 to 90°C (incremental steps of 0.5°C min^-1^). The melting curve was drawn using GraphPad Prism 7.0 software; the values of the first derivative of 350/330 nm were displayed on the Y axis. The melting temperature (Tm) was calculated as the mid-point temperature of the melting curve using the ThermControl software (NanoTemper Technologies).

#### Enzyme Assays

A fluorescent substrate harboring the cleavage site (indicated by ↓) of SARS CoV-2 Mpro (Dabcyl-KTSAVLQ↓SGFRKM-E(Edans)-NH_2_; GL Biochem) and buffer composed of 20 mM HEPES, 120 mM NaCl, 0.4 mM EDTA, 4 mM DTT, 20% glycerol, 0.5% DMSO, pH 7.0 was used for the inhibition assay. In the fluorescence resonance energy transfer (FRET)-based cleavage assay, the fluorescence signal of the Edans generated due to the cleavage of the substrate by the Mpro was monitored at an emission wavelength of 460 nm with excitation at 360 nm, using a Flx800 fluorescence spectrophotometer (BioTek). Initially, 10 *μ*L of SARS-CoV-2 Mpro WT at the final concentration of 50 nM, or SARS-CoV-2 Mpro H172Y at 400 nM, was pipetted into a 96-well plate containing pre-pipetted 60 *μ*L of reaction buffer. Subsequently, the reaction was initiated by addition of 30 *μ*L of the substrate dissolved in the reaction buffer to 100 *μ*L final volume, at different final concentrations varied from 10 to 320 *μ*M (10, 20, 40, 80, 120, 160, 240, 320 *μ*M). A calibration curve was generated by measurement of varied concentrations (from 0.04 to 6250 nM) of free Edans, with gain 80 in a final volume of 100 *μ*L reaction buffer. Initial velocities were determined from the linear section of the curve, and the corresponding relative fluorescence units per unit of time (ΔRFU/s) was converted to the amount of the cleaved substrate per unit of time (*μ*M/s) by fitting to the calibration curve of free Edans.

Inner-filter effect corrections were applied for the kinetic measurements according to Liu et al.^50^. The fluorescence of the substrate (in RFU) dissolved in 100 *μ*L final volume of reaction buffer at the corresponding concentrations used for the kinetic assay was measured and defined as f(substrate). Afterwards, 1 *μ*L free Edans was added (final concentration: 1 *μ*M) to each well, and the fluorescence reading was taken as f(substrate + Edans). Simultaneously, a reference value (in RFU) was measured with the same concentration of free Edans in 100 *μ*L of reaction buffer, giving f(reference). The inner-filter correction at each substrate concentration was calculated according to the function: corr % = (f (substrate + Edans) - f (substrate)) / f (reference) x 100%. The corrected initial velocity of the reaction was calculated as V = V_0_ / (corr%), where V_0_ represents the initial velocity of each reaction. As saturation could be achieved, kinetic constants (V_max_ and K_m_) were derived by fitting the corrected initial velocity to the Michaelis-Menten equation, *V* = *V*_max_ × [S]/(*K*_m_ + [S]), using GraphPad Prism 7.0 software. *k*_cat_/*K*_m_ was calculated according to the equation, *k*_cat_/*K*_m_ = *V*_max_/([*E*] × *K*_m_). Triplicate experiments were performed for each data point, and the value was presented as mean ± standard deviation (SD).

#### Determination of the IC_50_ of nirmatrelvir

The same substrate was employed as for the determination of the enzyme kinetics. The SPARK Multimode Microplate Reader (TECAN) was used to monitor the signal at same emission wavelength and excitation wavelength. The reaction buffer was 20 mM HEPES, 120 mM NaCl, 0.4 mM EDTA, 4 mM DTT, 20% glycerol, pH 7.0, to achieve a final concentration of 2% DMSO which is same as in the enzyme kinetics measurement. Stock solutions of the compounds were prepared with 100% DMSO. For the determination of the IC_50_, 50 nM of SARS-CoV-2 Mpro or 400 nM of SARS-CoV-2 Mpro H172Y was incubated with nirmatrelvir at various concentrations from 0 to 100 μM in reaction buffer at 37°C for 10 min. Afterwards, the FRET substrate at a final concentration of 10 *μ*M was added to each well, at a final total volume of 100 *μ*L, to initiate the reaction. The GraphPad Prism 7.0 software (GraphPad) was used for the calculation of the IC_50_ values. Measurements of inhibitory activity of nirmatrelvir were performed in triplicate and are presented as the mean±SD.

## Conflicts of interest

There are no conflicts to declare.

## Acknowledgement

V.M.O. and J.S. acknowledge National Institutes of Health (R01GM098818 and R01CA256557) for funding. The work of M.F.I., X.S., and R.H. was funded by the German Center for Infection Research (DZIF), project FF01.905.

## Data and Software Availability

The input files of all simulations as well as analysis and plotting scripts can be accessed at https://github.com/JanaShenLab/Mpro-_H172Y. Raw molecular dynamics trajectories can be obtained from the corresponding author upon request.

## Notes

### Competing Interest Statement

The authors have declared no competing interest.

### Summary of Updates

This version has been revised to include more methodologies, data and analysis.

