## Supporting Information for "An Integrative Approach to Dissect the Drug Resistance Mechanism of the H172Y Mutation of SARS-CoV-2 Main Protease"

### List of Tables

S1 Calculated p$K_a$'s of the WT and H172Y SARS-CoV-2 Mpros . . . . . . . . . S-4

S2 Summary of the computational studies performed in this work . . . . . . . . S-5

S3 Summary of relevant distances in the X-ray structures of WT and H172Y Mpros . . . . . . . . . . . . . . . . . . . . . . . . . . . . . . . . . . . . . . S-6

S4 Five distances that most correlate with the frame label (i.e. apo or holo) for both Diffnet models. Pairwise distances were calculated between alpha carbons within 15 Åof the alpha carbon of H/Y172, using the centroid frame of each cluster (200 total) after clustering on the latent space. . . . . . . . . S-6

### List of Figures

S1 The overall RMSD of H172Y Mpro in all runs of the free and nirmatrelvir-bound H172Y Mpro . . . . . . . . . . . . . . . . . . . . . . . . . . . . . S-7

S2 Nirmatrelvir was stably bound in the WT and H172Y Mpros during the simulations. . . . . . . . . . . . . . . . . . . . . . . . . . . . . . . . . . . . . S-8

S3 Probability distributions of the relevant distances involving the S1 pocket in the free WT Mpro . . . . . . . . . . . . . . . . . . . . . . . . . . . . . . S-9

S4 Run 1 of the free H172Y Mpro . . . . . . . . . . . . . . . . . . . . . . . . S-10

S5 Run 2 of the free H172Y Mpro . . . . . . . . . . . . . . . . . . . . . . . . S-11

S6 Run 3 of the free H172Y Mpro . . . . . . . . . . . . . . . . . . . . . . . . S-12

S7 **H172Y causes a shift between His163 and Phe140 that can disrupt stacking.** Probability distributions for the distance between the COM of aromatic rings of Phe140 and His163 for ligand-free trajectories after removing the first 1 $\mu$. The distribution from each trajectory is shown as separate lines, colored according to the system (WT or mutant), separated by monomer. . . . . . . . . . . . . . . . . . . . . . . . . . . . . . . . . S-13
S8 Stability of the oxyanion loop in the simulations of the free and nirmatrelvir-bound H172Y Mpro . . . . . . . . . . . . . . . . . . . . . . . . . . . . S-14
S9 Occupancy of the nonnative hydrogen bond between Phe140 and Tyr172 in run 1 of the ligand-free H172Y Mpro . . . . . . . . . . . . . . . . . . . . . . S-14
S10 Possible correlation between the formation of the Phe140–Tyr172 nonnative hydrogen bond and the loss of aromatic stacking . . . . . . . . . . . . . S-15
S11 Run 1 of the nirmatrelvir bound H172Y Mpro . . . . . . . . . . . . . . . . . . S-16
S12 Run 2 of the nirmatrelvir bound H172Y Mpro . . . . . . . . . . . . . . . . . . S-17
S13 Nirmatrelvir binding interactions are affected by the H172Y mutation . . . . S-18
S14 Performance of the autoencoder. . . . . . . . . . . . . . . . . . . . . . . . . S-19
S15 H172Y induces conformational change to the oxyanion loop in both the apo and holo trajectories . . . . . . . . . . . . . . . . . . . . . . . . . . . . . . S-20
S16 H172Y induces conformational changes to the region preceding the oxyanion loop in both apo and holo trajectories. . . . . . . . . . . . . . . . . . . . S-21

### Supplemental Tables

Table S1: Summary of the p$K_a$'s of the WT and H172Y mutant SARS-CoV-2 Mpros calculated from the CpHMD titration simulations[a]

| | WT Mpro | | H172Y Mpro | |
|---|---|---|---|---|
| Residue | 6y2g(A) | 6y2g(B) | 7vh8(A) | 7vh8(B) |
| H41 | 6.6 | 6.7 | 6.6 | 6.2 |
| H64 | 6.1 | 6.1 | 6.2 | 6.1 |
| H80 | 6.3 | 6.2 | 6.1 | 6.0 |
| H163 | neutral[b] | neutral | neutral | neutral |
| H164 | neutral | neutral | neutral | neutral |
| H172 | 6.6 | 6.6 | N/A[c] | N/A |
| H246 | ∼4.3[d] | ∼4.9 | 5.4 | 5.0 |
| C16 | neutral | neutral | neutral | neutral |
| C22 | 7.5 | 6.8 | neutral | ∼7.3 |
| C38 | neutral | neutral | neutral | neutral |
| C44 | 7.0 | 9.2 | ∼7.1 | ∼7.7 |
| C85 | ∼9.5 | ∼9.0 | ∼9.2 | neutral |
| C117 | neutral | ∼9.2 | neutral | neutral |
| C128 | ∼9.4 | ∼9.1 | ∼9.4 | ∼8.9 |
| C145 | neutral | ∼9.4 | neutral | ∼9.3 |
| C156 | neutral | neutral | ∼8.3 | ∼8.8 |
| C160 | neutral | neutral | neutral | neutral |
| C265 | neutral | neutral | neutral | neutral |
| C300 | 8.4 | neutral | neutral | neutral |

[a] The p$K_a$'s of both protomers are listed here. The results of the WT Mpro are taken from our previous paper.[S1] Simulations of the H172Y mutant were initiated using the computationally mutated H172 mutant structure based on the X-ray structure of the WT Mpro (PDB id 7vh8,[S2] ligand removed). The entire simulation time (30 ns per pH replica) was used for calculation of the protonation fractions. [b] For histidines, neutral indicates that the residue remains in the singly protonated state (i.e., charge neutral) in the entire simulation pH range. For cysteines, neutral indicates that the residue remains in the protonated state (i.e., charge neutral) in the entire simulation pH range. [c] N/A indicates H172 does not exist. [d] ∼ indicates that due to the incomplete titration (in the simulation pH range) the calculated p$K_a$ is approximate.

Table S2: Summary of the computational studies performed in this work

| Protein | PDB ID | Set up | Simulation time |
|---|---|---|---|
| **Replica-exchange CpHMD**[S3] | | | |
| H172Y | modeled[a] | ligand free | 30 ns x 9 |
| **Fixed-protonation state MD**[S4] | | | |
| WT | 7vh8 | ligand free | 2 $\mu$s x 3 |
| WT | 7vh8 | nirmatrelvir bound[b] | 2 $\mu$s x 2 |
| H172Y | modeled[a] | ligand free | 2 $\mu$s x 3 |
| H172Y | unpublished (Hilgenfeld group) | ligand-free | 2 $\mu$s x 1 |
| H172Y | modeled[a] | nirmatrelvir bound[b] | 2 $\mu$s x 2 |
| **Alchemical free energy perturbation**[S5–S7] | | | |
| WT$\rightarrow$H172Y | 7vh8[c] | nirmatrelvir bound[b] | 24 ns[e] $\times$ 2 |
| WT$\rightarrow$H172Y | 7vh8[c] | ligand free[d] | 24 ns[e] $\times$ 2 |
| H172Y$\rightarrow$WT | modeled[d] | nirmatrelvir bound[b] | 24 ns[e] $\times$ 2 |
| H172Y$\rightarrow$WT | modeled[d] | ligand free | 24 ns[e] $\times$ 2 |

| Protein | PDB ID | Set up | Num. of repeats |
|---|---|---|---|
| **Empirical protein stability calculations with Rosetta ddG_monomer**[S8] | | | |
| WT | 7vh8 | ligand free | 50 |
| H172Y | modeled[f] | ligand free | 50 |
| **Empirical binding free energy calculations with Rosetta flex ddG**[S9] | | | |
| WT$\rightarrow$ H172Y[g] | 7vh8 | free or ligand bound | 40 |
| H172Y$\rightarrow$WT[g] | modeled[a] | free or ligand bound | 40 |

[a]The H172Y mutation was introduced using Modeller[S10] to the X-ray structure of the WT Mpro in complex with nirmatrelvir (PDB id 7vh8)[S2] as the template. [b]The PDB entry 7vh8[S2] contains the structures of WT Mpro-nirmatrelvir complex in both the covalent and noncovalent binding modes. The latter was used as the starting structure of the simulations. [c]The dual topology hybrid molecule (H172/Y172) was created following 50-ns equilibration simulation of the WT or H172Y Mpro. [d]Mutant structure was built using VMD[S11] based on the WT Mpro structure (PDB id 7vh8).[S2] [e]Sampling time includes the 12-ns forward and backward transformations. [f]Generated by the Rosetta ddG_monomer program.[S8] [g]Mutation introduced by the Rosetta flex ddG program.[S9]

Table S3: Summary of relevant distances in the X-ray structure of WT Mpro and in the newly reported X-ray structures of the H172Y Mpro[a]

| Mpro<br>Ligand | 7vh8 (WT)[S2]<br>nirmatrelvir | 8d4j (H172Y)[S12]<br>free | 8d4k (H172Y)[S12]<br>GC-376 |
|---|---|---|---|
| F140–H163 | 3.7 | 3.8/3.8 | 3.7/3.7 |
| F140:N–Y172:OH | n/a | 3.5/3.6 | 3.3/3.3 |
| Y172:OH–S1*:N | n/a | 4.1/4.4 | n/d |
| H172:ND1/NE2–S1*:O | 3.1 | n/a | n/a |
| E166:OE1/2–S1*:N | 2.8 | 4.3/3.3 | n/d |
| F140:O–S1*:N | 2.8 | 6.5/6.5 | n/d |
| G138:CA–S144:CA | 10.9 | 11.1 | 11.1 |
| G138:CA–T135:CA | 9.0 | 9.2 | 9.5 |

[a]All distances are in unit Å. F140–H163 refers to the distance between the center-of-mass (COM) of the aromatic rings of Phe140 and His163. The PDB entry 7vh8, the WT Mpro is in complex with nirmatrelvir. In the PDB entry 8d4k, the coordinates of Ser1 are not resolved; thus, distances involving S1 are listed as n/d. 8d4j: ligand free H172Y Mpro. 8d4k: H172Y Mpro in complex with a covalent inhibitor GC-376.

Table S4: Five distances that most correlate with the frame label (i.e. apo or holo) for both Diffnet models. Pairwise distances were calculated between alpha carbons within 15 Åof the alpha carbon of H/Y172, using the centroid frame of each cluster (200 total) after clustering on the latent space.

Apo

| Residue:Chain | Residue:Chain | Correlation w/ predicted label |
|---|---|---|
| Gly138:B | Thr135:B | 0.9043 |
| Gly138:B | Asn133:B | 0.8756 |
| Gly138:B | Phe134:B | 0.8661 |
| Gln127:B | Ala129:B | 0.8427 |
| Phe181:B | Phe3:B | 0.8356 |

Holo

| | | |
|---|---|---|
| Gly138:A | Gly143:A | 0.9257 |
| Gly138:A | Ser144:A | 0.9073 |
| Glu166:A | Ala173:A | 0.9017 |
| Asn142:A | Gly138:A | 0.8802 |
| Phe112:B | Phe3:B | 0.8700 |

### Supplemental Figures

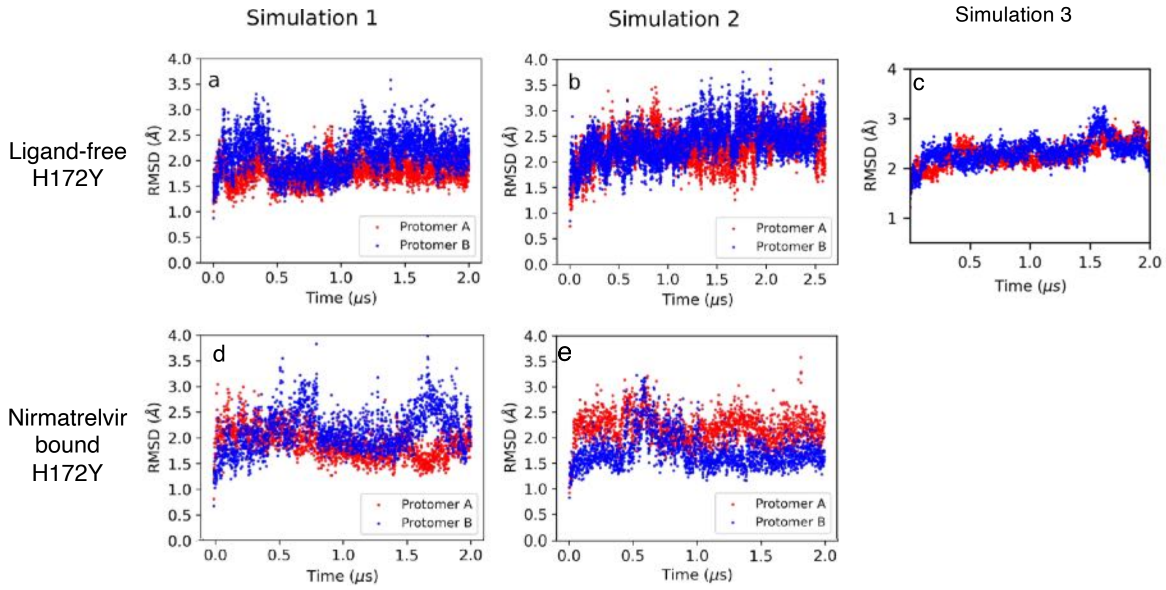


Figure S1: **The overall RMSD of H172Y Mpro in Run 1 and 2 of the free and nirmatrelvir-bound H172Y Mpro.** Heavy-atom root-mean-square deviation (RMSD) of the ligand-free (a, b, c) and nirmatrelvir bound (d, e) H172Y Mpros with respect to the mutant model as a function of simulation time. Simulation runs 1 and 2 are shown on the left and center panels, respectively, while the third free simulation is shown on the right.

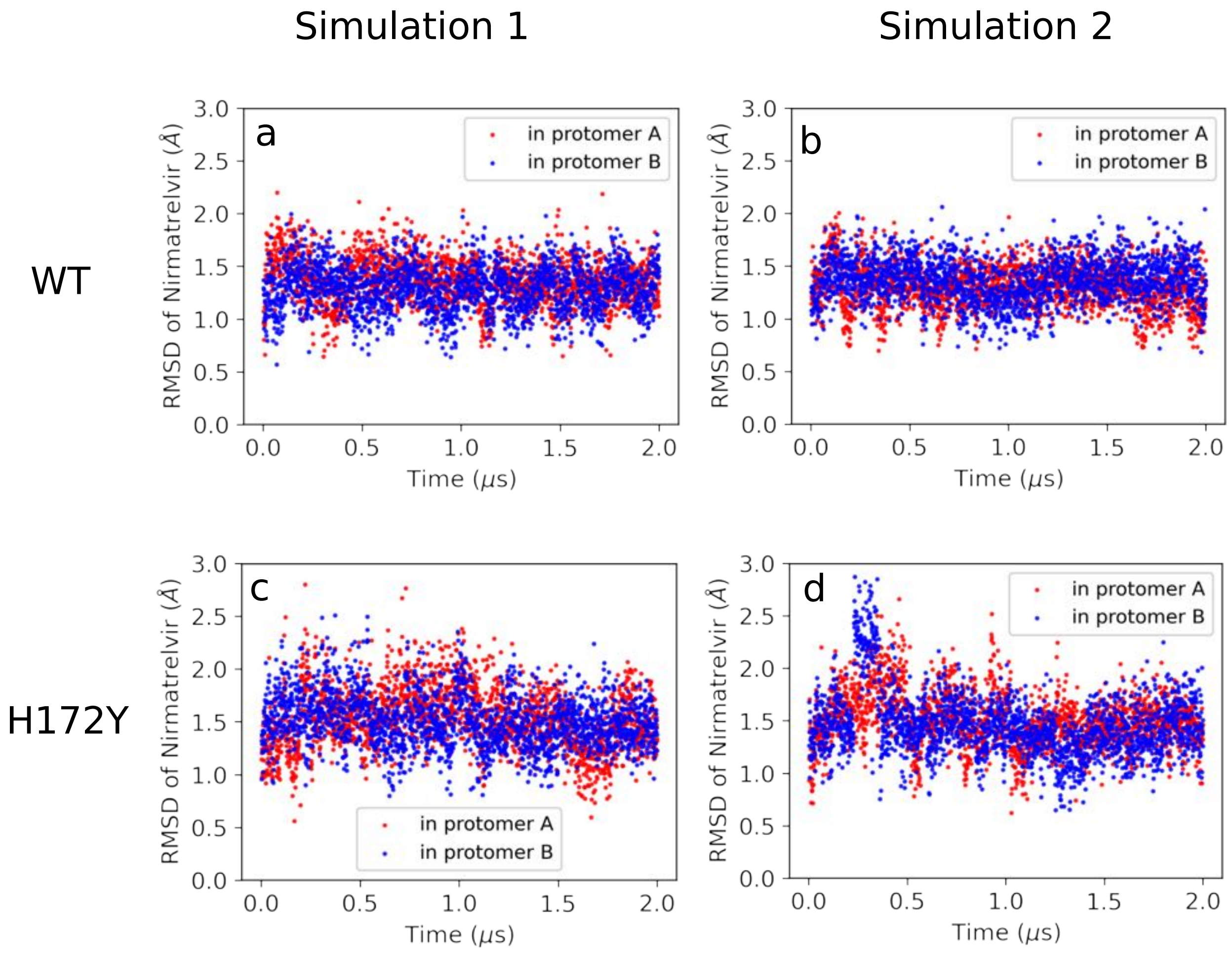


Figure S2: **Nirmatrelvir was stably bound in the WT and H172Y Mpros during the simulations.** (a, b) Time series of the RMSD of nirmatrelvir in the WT Mpro with respect to the crystal structure (PDB id 7vh8) in the simulation run 1 (a) and 2 (b). (c, d) Time series of the RMSD of nirmatrelvir in the H172Y Mpro with respect to the mutant model in the simulation run 1 (c) and 2 (d).

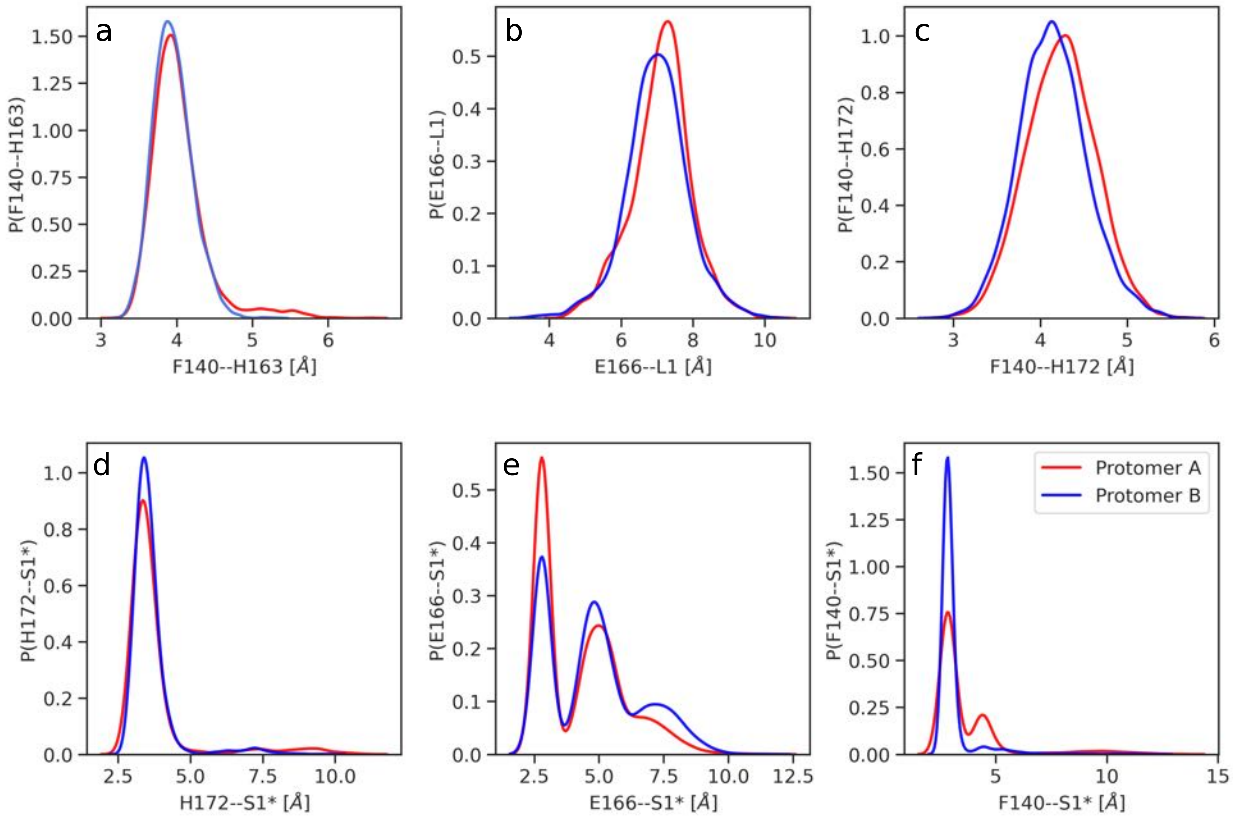


Figure S3: **Probability distributions of the distances involving the S1 pocket residues in the free WT Mpro.** Distributions of the distance (a) between the center-of-mass (COM) of the aromatic rings of Phe140 and His163; (b) between the COM of the carboxylate oxygens of Glu166 and C$\alpha$ atoms of residues L1; (c) between the amide nitrogen of Phe140 and the hydroxyl oxygen of His172; (d) between the backbone carbonyl oxygen of S1$^*$ and the nearest imidazole nitrogen of His172; (e) between the amino nitrogen of Ser1$^*$ and the nearest carboxylate oxygen of Glu166. (f) between the backbone carbonyl oxygen of Phe140 and the amino nitrogen of Ser1$^*$. The red and blue curves represent protomer A and B, respectively. The calculations used the data from both simulation runs of the free WT Mpro. The most probable (peak) distances were used as references in the analysis of the H172Y Mpro simulations.

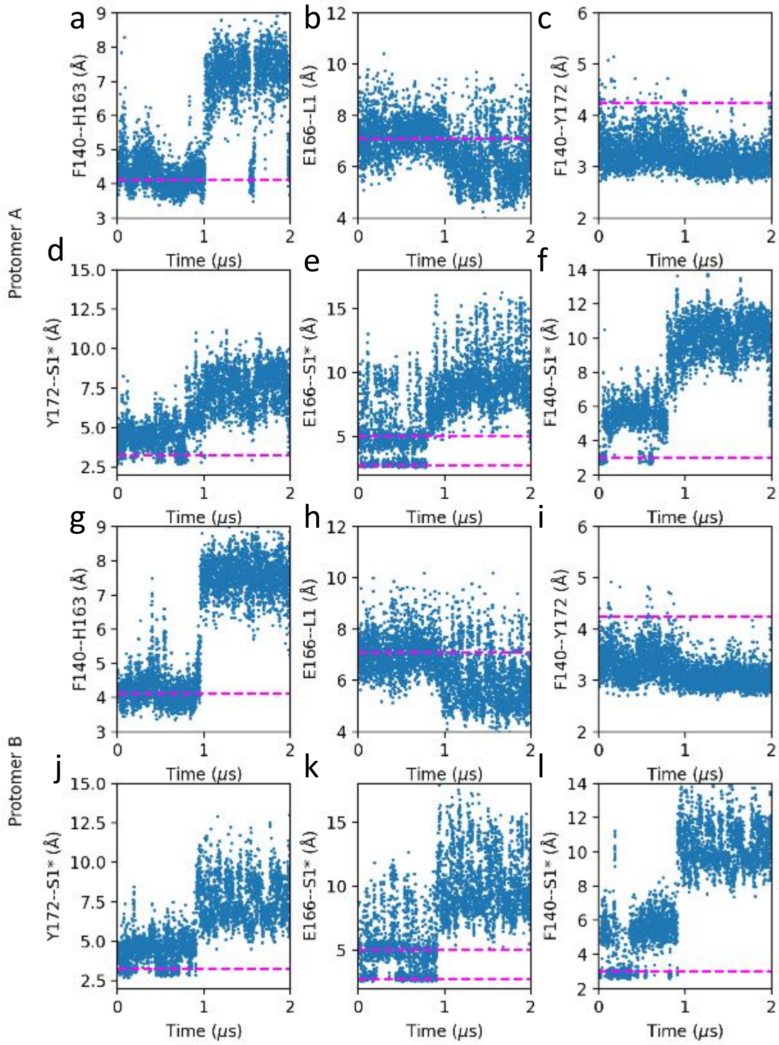


Figure S4: **Run 1 of the ligand-free H172Y Mpro: the S1 pocket–Ser1$^*$ interactions and the Phe140–His163 stacking are disrupted.** Distributions of the distances between the center-of-mass (COM) of the aromatic rings of Phe140 and His163 (a); between the COM of the carboxylate oxygens of Glu166 and C$\alpha$ atoms of L1 (b); between the amide nitrogen of Phe140 and the hydroxyl oxygen of Tyr172 (c); between the amino nitrogen of Ser1$^*$ and the hydroxyl group of Tyr172 (d); the nearest carboxylate oxygen of Glu166 (e); or the backbone carbonyl oxygen of Phe140 (f). The magenta lines represent the most probable distances sampled by the free WT Mpro.

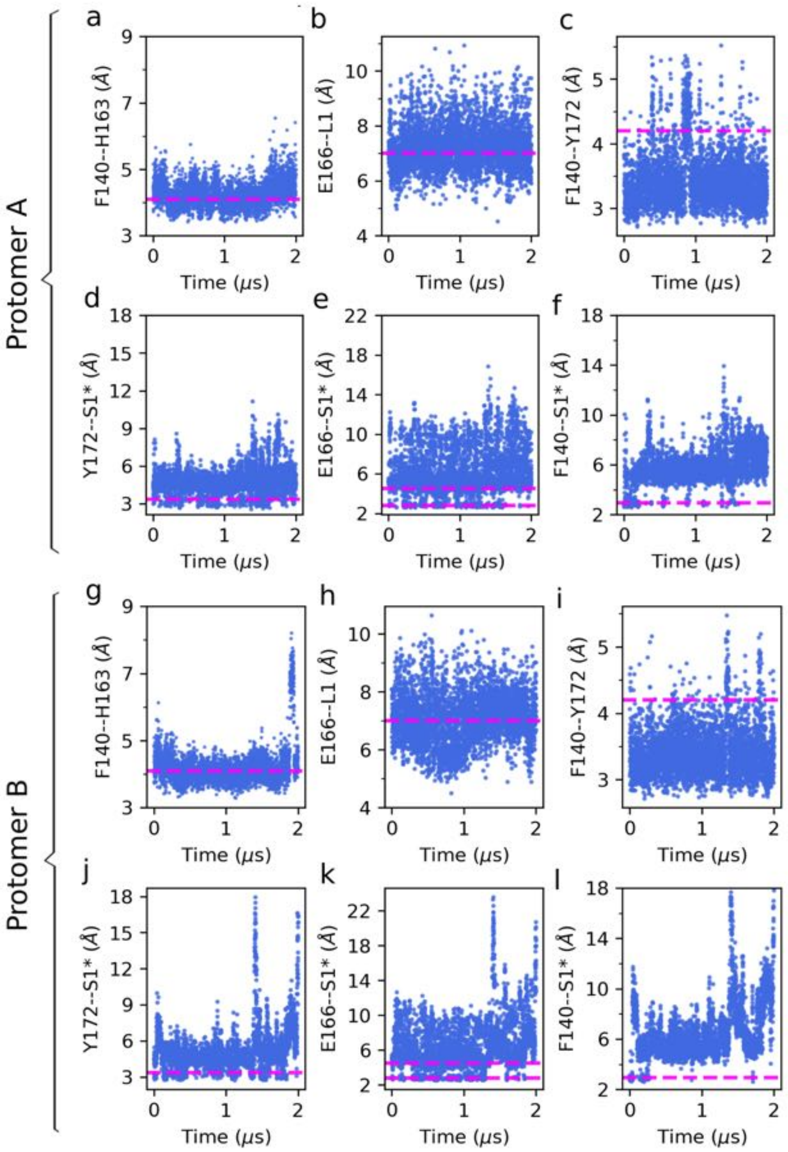


Figure S5: **Run 2 of the free H172Y Mpro: the S1 pocket–Ser1$^*$ interactions are disrupted and the Phe140–His163 is unstable in both protomers.** Distributions of the distances between the center-of-mass (COM) of the aromatic rings of Phe140 and His163 (a,g); between the COM of the carboxylate oxygens of Glu166 and C$\alpha$ atoms of L1 (b,h); between the amide nitrogen of Phe140 and the hydroxyl oxygen of Tyr172 (c,i); between the amino nitrogen of Ser1$^*$ and the hydroxyl group of Tyr172 (d,j); the nearest carboxylate oxygen of Glu166 (e,k); or the backbone carbonyl oxygen of Phe140 (f,i). The magenta lines represent the most probable distances sampled by the free WT Mpro.

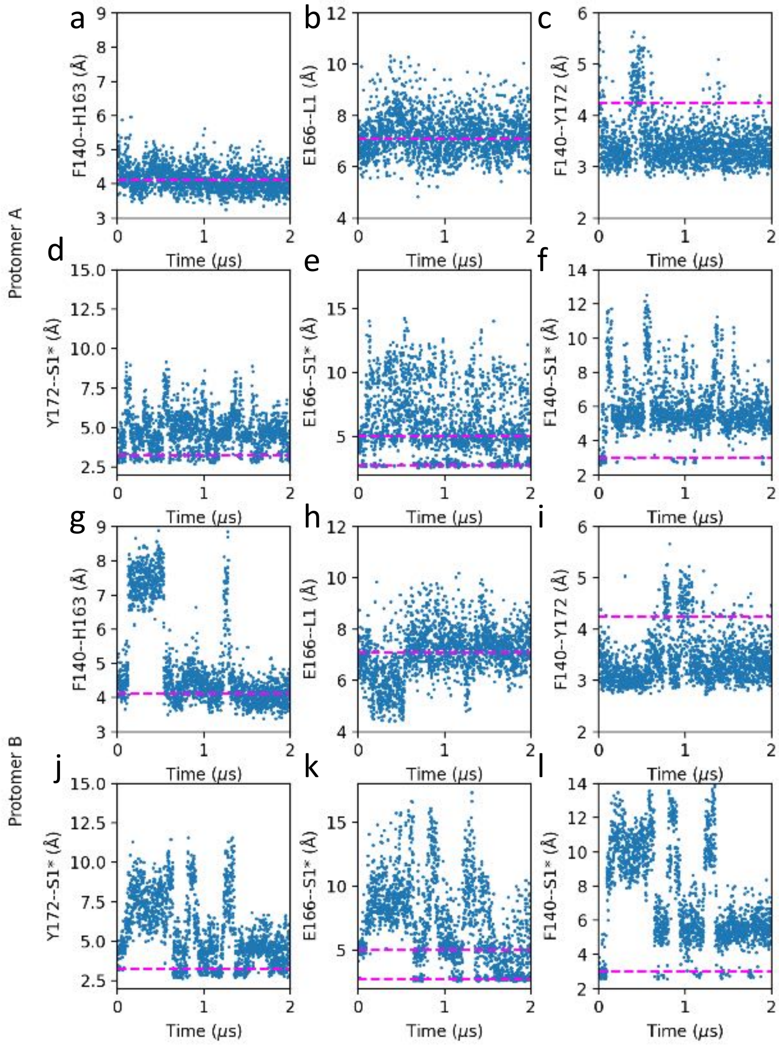


Figure S6: **Run 3 of the free H172Y Mpro: the S1 pocket–Ser1$^*$ interactions are disrupted and the His163-Phe140 stacking in protomer B is unstable.** Distributions of the distances between the center-of-mass (COM) of the aromatic rings of Phe140 and His163 (a,g); between the COM of the carboxylate oxygens of Glu166 and C$\alpha$ atoms of L1 (b,h); between the amide nitrogen of Phe140 and the hydroxyl oxygen of Tyr172 (c,i); between the amino nitrogen of Ser1$^*$ and the hydroxyl group of Tyr172 (d,j); the nearest carboxylate oxygen of Glu166 (e,k); or the backbone carbonyl oxygen of Phe140 (f,i).

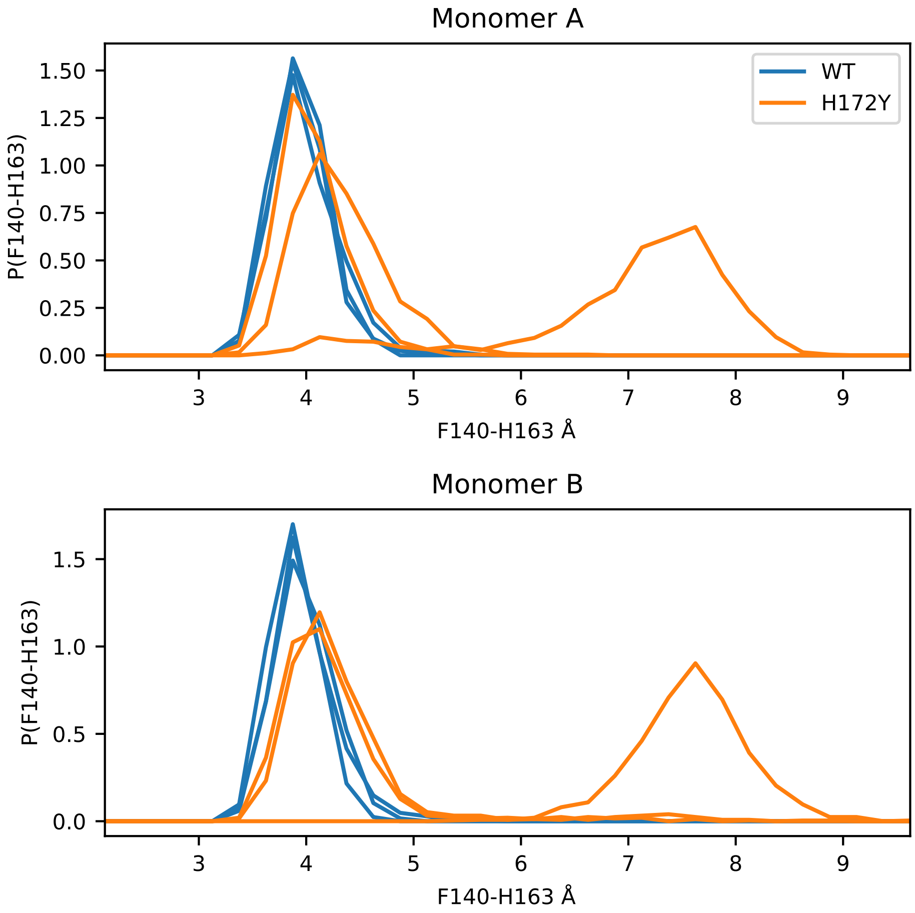


Figure S7: **H172Y causes a shift between His163 and Phe140 that can disrupt stacking.** Probability distributions for the distance between the COM of aromatic rings of Phe140 and His163 for ligand-free trajectories after removing the first 1 $\mu$. The distribution from each trajectory is shown as separate lines, colored according to the system (WT or mutant), separated by monomer.

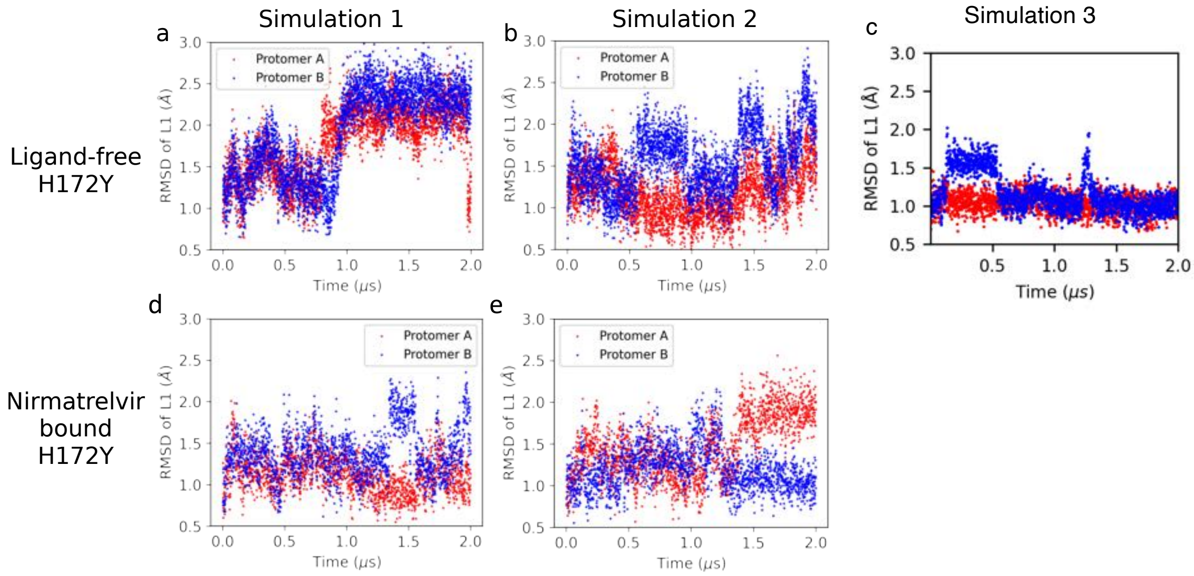


Figure S8: **Stability of the oxyanion loop in the simulations of the free and nirmatrelvir-bound H172Y Mpro.** Heavy-atom RMSD of the Mpro oxyanion loop (residues 138-145, L1) with respect to the to the mutant model as a function of simulation time in the ligand-free (a, b) and nirmatrelvir bound H172Y (c, d) Mpros. Simulation runs 1 and 2 are shown on the left and right panels, respectively. Run 3 is shown on the bottom.

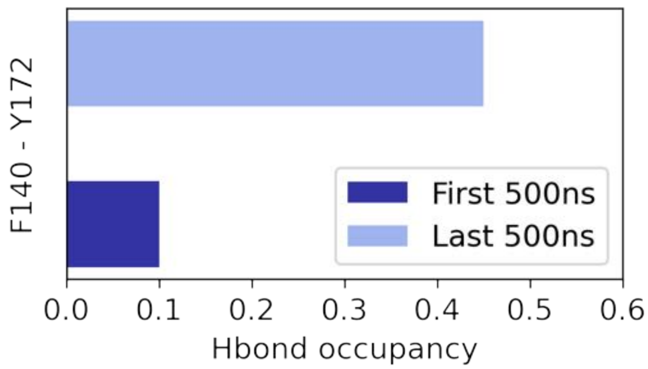


Figure S9: **Run 1 of the ligand-free H172Y Mpro: Occupancy of the nonnative hydrogen bond between amide nitrogen of Phe140 and the hydroxyl oxygen of Tyr172.** The occupancy was calculated for the first 0.5 $\mu$s (dark blue) and last 0.5 $\mu$s of the 2-$\mu$s simulation.

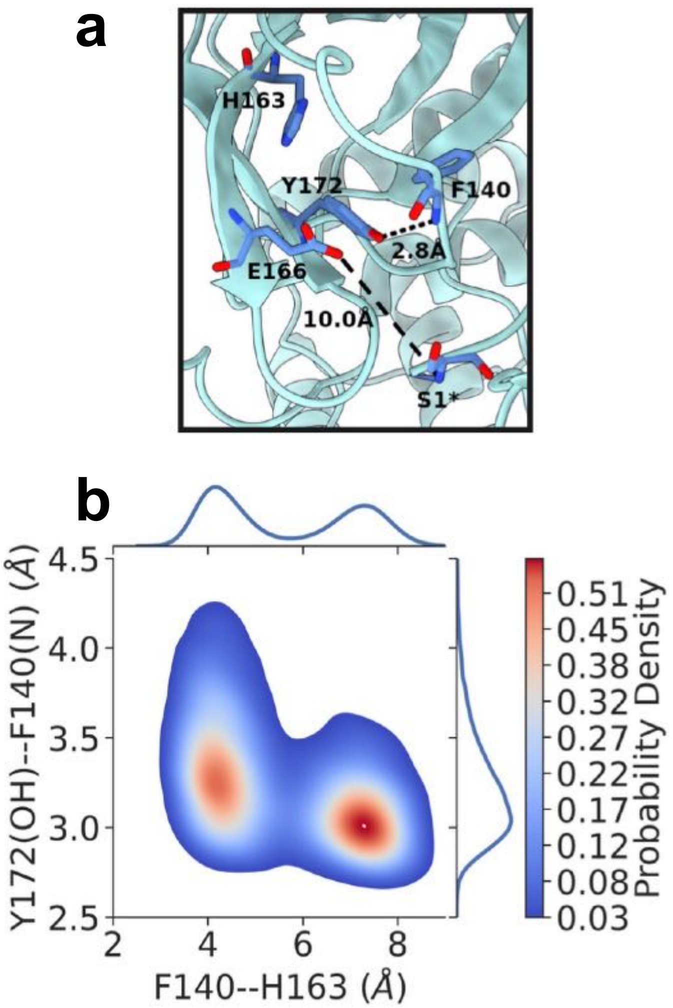


Figure S10: **Formation of the nonnative Y172-F140 hydrogen bond may be correlated with the loss of aromatic stacking in the free H172Y Mpro. a.** A representative structure (cluster centroid) was taken from the clustering analysis of the last 1-$\mu$s trajectory. A zoomed-in view of the S1 pocket shows a nonnative hydrogen bond between the hydroxyl group of Tyr172 and the backbone amide of Phe140, the complete loss of aromatic stacking between His163 and Phe140 and the N terminus interaction with Glu166. The Y172:O–F140:N and E166:OE1/OE2–S1$^*$:N distances are indicated. **b.** Probability density as a function of the F140–H163 and F140–Y172 distances from the last 1-$\mu$s trajectory. Analysis here uses the data of protomer A from the simulation run 1 of the free H172Y Mpro simulations. **b.**

dfd

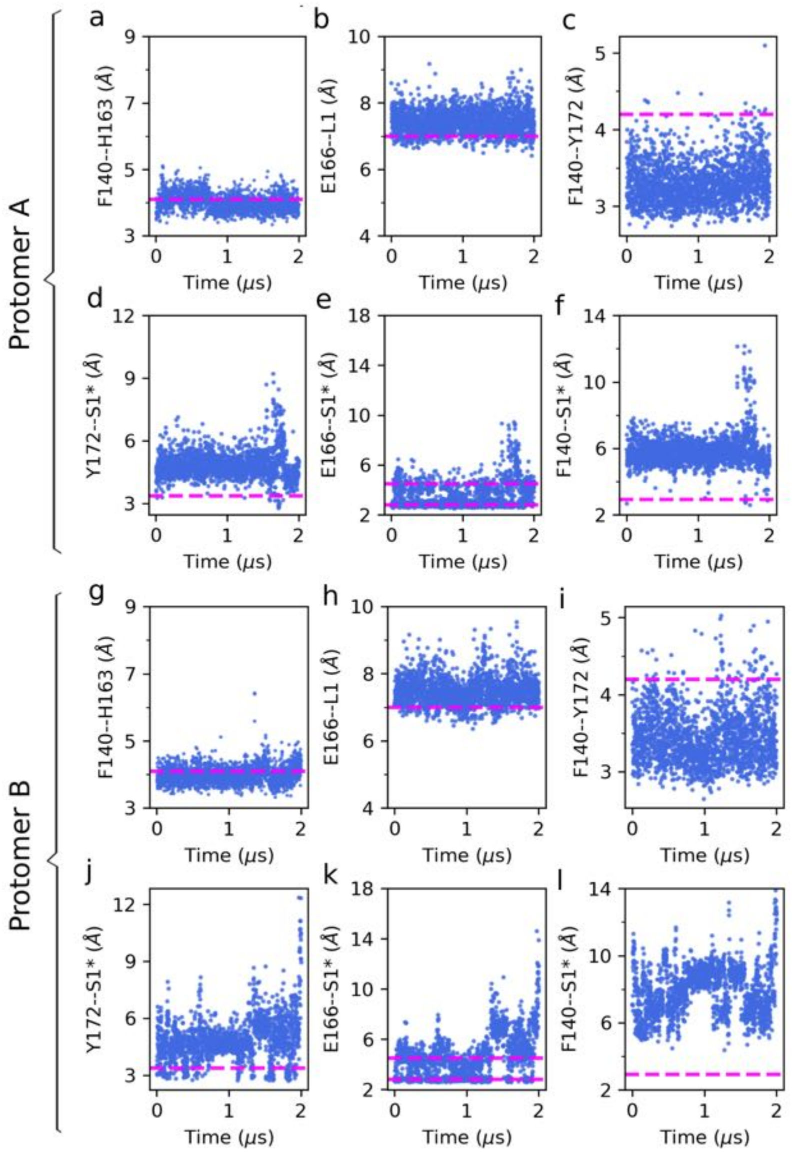


Figure S11: **Run 1 of the nirmatrelvir bound H172Y Mpro: the S1 pocket–Ser1$^*$ interactions were disrupted.** (a, g) Distance between the COM of the aromatic rings of Phe140 and His163 in protomer A (a) and B (g). (b, h) Distance between the center of mass (COM) of the carboxylate oxygens of Glu166 and that of the oxyanion loop (C$\alpha$ atoms of residues 138-145) in protomer A (b) and B (h). (c, i) Distance between the amide nitrogen of Phe140 and the hydroxyl oxygen of Tyr172 in protomer A (c) and B (i). (d, j) Distance between the hydroxyl group of Tyr172 and the N-terminus amino nitrogen in the opposite protomer (Ser1*). (e, k) Distance between the nearest carboxylate oxygen of Glu166B and the amino nitrogen of Ser1$^*$. (f, l) Distance between the backbone carbonyl oxygen of Phe140 and the amino nitrogen of Ser1$^*$. The magenta dashed lines in the plots represent the average distances sampled by WT simulations.

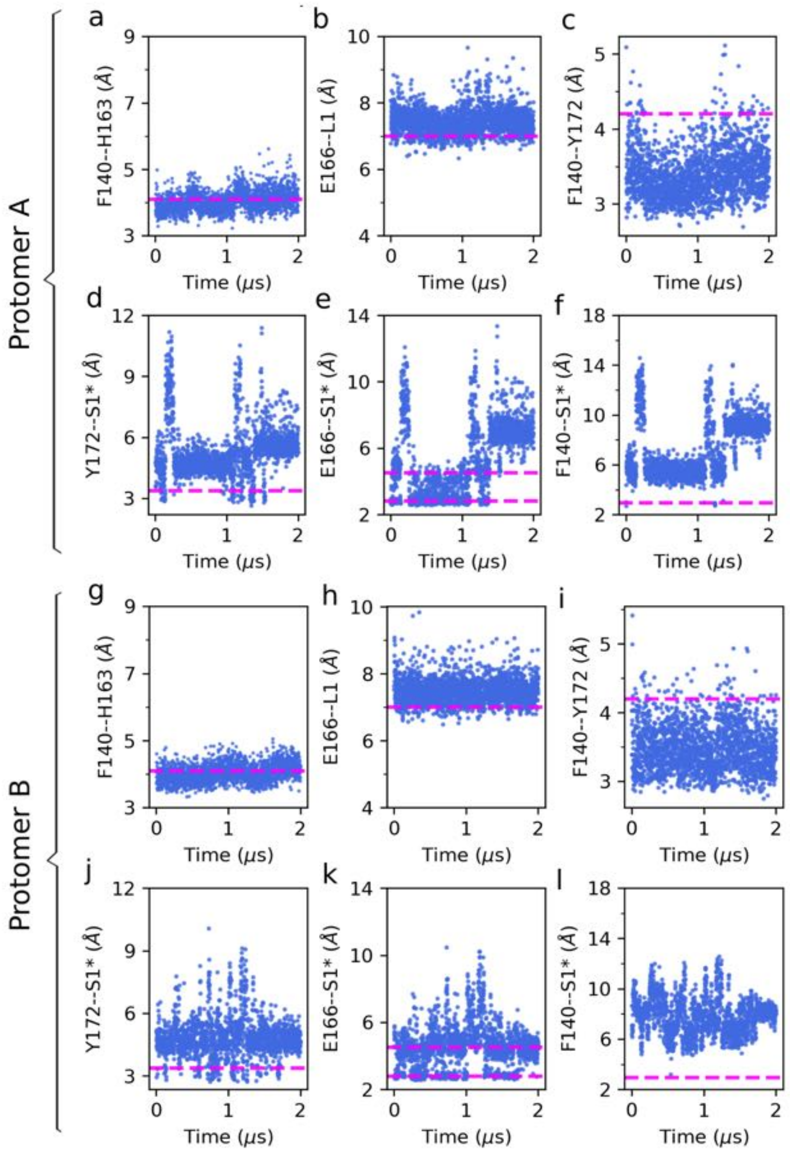


Figure S12: **Run 2 of the nirmatrelvir bound H172Y Mpro: the S1 pocket – Ser1* interactions were disrupted.** (a, g) Distance between the COM of the aromatic rings of Phe140 and His163 in protomer A (a) and B (g). (b, h) Distance between the center of mass (COM) of the carboxylate oxygens of Glu166 and that of the oxyanion loop (C$\alpha$ atoms of residues 138–145) in protomer A (b) and B (h). (c, i) Distance between the amide nitrogen of Phe140 and the hydroxyl oxygen of Tyr172 in protomer A (c) and B (i). (d, j) Distance between the hydroxyl group of Tyr172 and the amino nitrogen of Ser1*. (e, k) Distance between the nearest carboxylate oxygen of Glu166B and the amino nitrogen of Ser1*. (f, l) Distance between the backbone carbonyl oxygen of Phe140 and the amino nitrogen of Ser1*. The magenta dashed lines in the plots represent the average distances sampled by WT simulations.

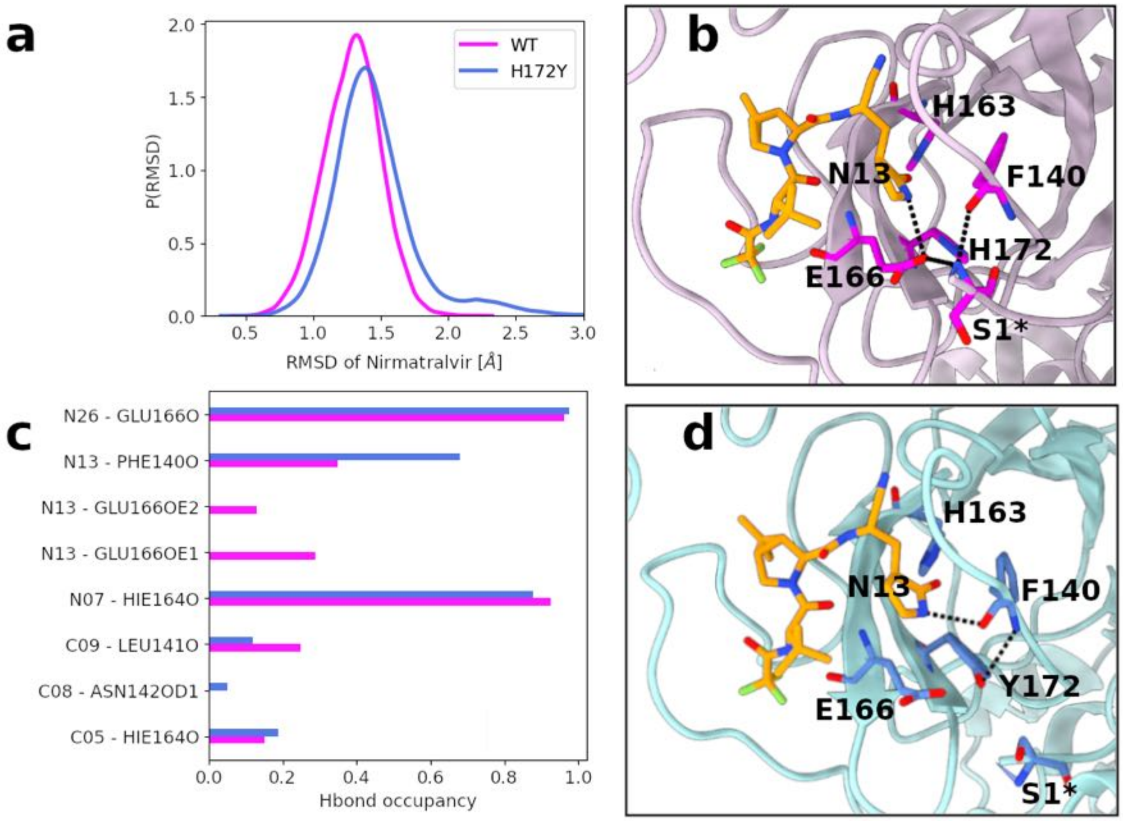


Figure S13: **Binding interactions between nirmatrelvir and Mpro are affected by the H172Y mutation.** (a) Probability distributions of the heavy atom RMSD of nirmatrelvir in the WT (magenta) and H172Y (blue) Mpros with respect to the X-ray structure (PDB id 7vh8). Data from both simulation runs were used. b) A zoomed-in view of the nirmatrelvir binding site in the WT Mpro. (c) Occupancies of the hydrogen bonds between nirmatrelvir and Mpro atoms in the WT (magenta) and H172Y (blue) Mpros. The data from the last 1 $\mu$s of both simulation runs were used. (d) A zoomed-in view of the nirmatrelvir binding site in the H172Y Mpro based on the representative (clustering centroid) structure. A nonnative hydrogen bond is formed between the backbone of Phe140 and sidechain of Tyr172. The hydrogen bond betwen the lactam N13 and Glu166 carboxylate oxygen is missing. Instead, a hydrogen bond between N13 and the backbone carbonyl of Phe140 is formed.

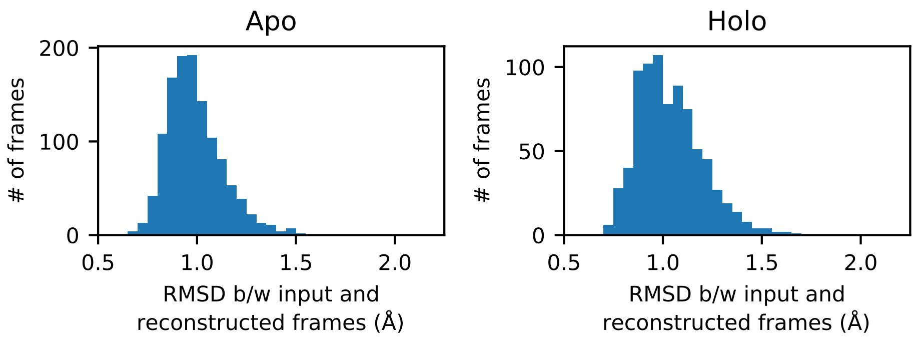


Figure S14: **Created diffnets are capable of reproducing the input frame after encoding onto the latent space.** Performance evaluation of diffnets trained on apo (left) and holo (right) trajectories. The RMSD between the input (i.e. frame from an MD trajectory) and the reconstructed frames (i.e. frame created by the decoder). Both models are adept at reconstructing the input positions, with RMSD between input and reconstruction ranging from 0.6 and 1.7 Å.

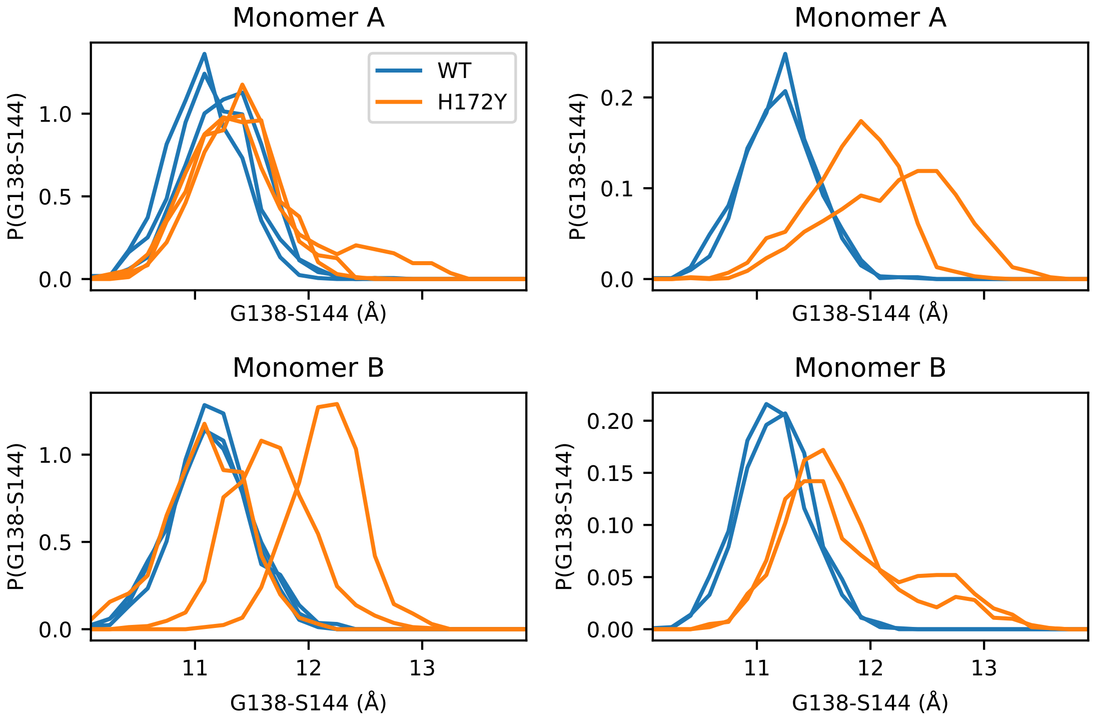


Figure S15: **H172Y induces conformational changes to the oxyanion loop in both the apo and holo trajectories.** Probability distributions for the distance between C$\alpha$ atoms of G138 and S144 (one residue before the last residue in the oxyanion loop). The distribution from each trajectory is shown as a separate line, colored according to WT or mutant, separated by system (apo or holo) and by monomer.

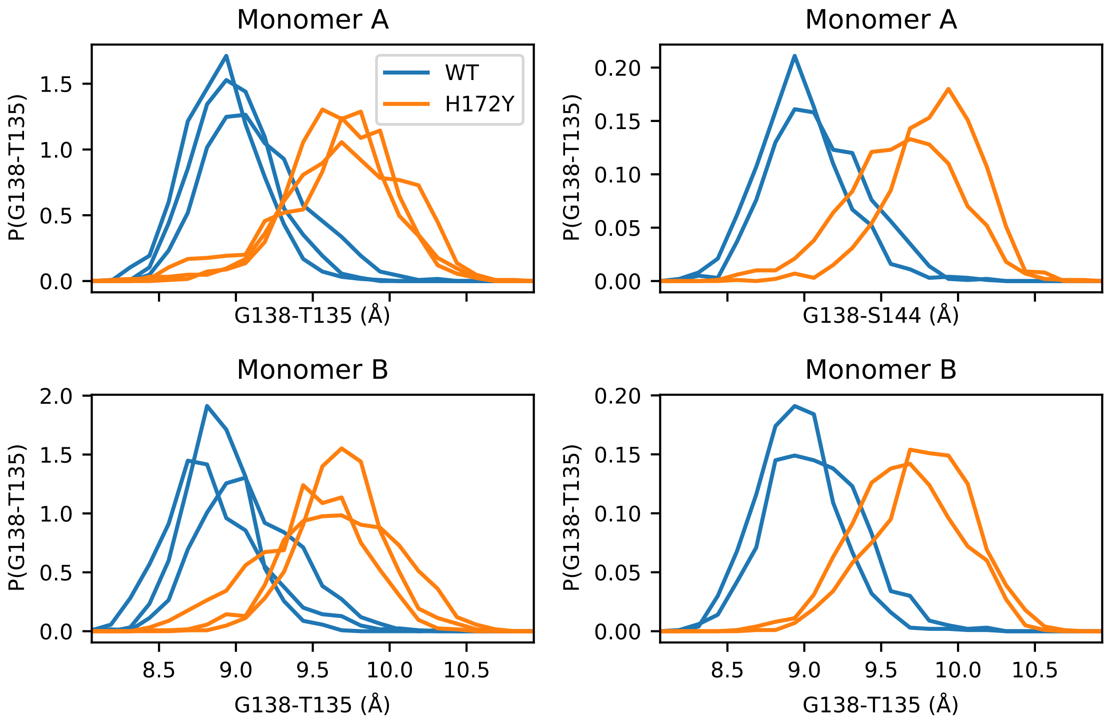


Figure S16: **H172Y induces conformational changes to the region preceding the oxyanion loop in both apo and holo trajectories.** Probability distributions for the distance between alpha carbons of residues preceding the oxyanion loop, G138 and T135. The distribution from each trajectory is shown as a separate line, colored according to WT or mutation, separated by system (apo or holo) and by monomer.

(S4) Case, D. A.; Ben-Shalom, I. Y.; Brozell, S. R.; Cerutti, D. S.; Cheatham, T., III; Cruzeiro, V. W. D.; Darden, T. A.; Duke, R. E.; Ghoreishi, D.; Gilson, M. K.; Gohlke, H.; Goetz, A. W.; Greene, D.; Harris, R.; Homeyer, N.; Huang, Y.; Izadi, S.; Kovalenko, A.; Kurtzman, T.; Lee, T. S.; LeGrand, S.; Li, P.; Lin, C.; Liu, J.; Luchko, T.; Luo, R.; Mermelstein, D. J.; Merz, K. M.; Miao, Y.; Monard, G.; Nguyen, C.; Nguyen, H.; Omelyan, I.; Onufriev, A.; Pan, F.; Qi, R.; Roe, D. R.; Roitberg, A.; Sagui, C.; Schott-Verdugo, S.; Shen, J.; Simmerling, C. L.; Smith, J.; Salomon-Ferrer, R.; Swails, J.; Walker, R. C.; Wang, J.; Wei, H.; Wolf, R. M.; Wu, X.; Xiao, L.; York, D. M.; Kollman, P. A. AMBER 2020. 2020.

(S5) Dixit, S. B.; Chipot, C. Can Absolute Free Energies of Association Be Estimated from Molecular Mechanical Simulations? The BiotinStreptavidin System Revisited. *The Journal of Physical Chemistry A* **2001**, *105*, 9795–9799.

(S6) Phillips, J. C.; Braun, R.; Wang, W.; Gumbart, J.; Tajkhorshid, E.; Villa, E.;

Chipot, C.; Skeel, R. D.; Kalé, L.; Schulten, K. Scalable Molecular Dynamics with NAMD. *J Comput Chem* **2005**, *26*, 1781–1802.

(S7) Jiang, W.; Chipot, C.; Roux, B. Computing Relative Binding Affinity of Ligands to Receptor: An Effective Hybrid Single-Dual-Topology Free-Energy Perturbation Approach in NAMD. *J. Chem. Inf. Model.* **2019**, *59*, 3794–3802.

(S8) Kellogg, E. H.; Leaver-Fay, A.; Baker, D. Role of conformational sampling in computing mutation-induced changes in protein structure and stability. *Proteins: Structure, Function, and Bioinformatics* **2011**, *79*, 830–838.

(S9) Barlow, K. A.; Ó Conchúir, S.; Thompson, S.; Suresh, P.; Lucas, J. E.; Heinonen, M.; Kortemme, T. Flex ddG: Rosetta Ensemble-Based Estimation of Changes in Protein–Protein Binding Affinity upon Mutation. *J. Phys. Chem. B* **2018**, *122*, 5389–5399.

(S10) Webb, B.; Sali, A. Comparative Protein Structure Modeling Using MODELLER. *Current Protocols in Bioinformatics* **2016**, *54*, 5.6.1–5.6.37.

(S11) Humphrey, W.; Dalke, A.; Schulten, K. VMD: Visual Molecular Dynamics. *J Mol Graph* **1996**, *14*, 33–38.

(S12) Hu, Y.; Lewandowski, E. M.; Tan, H.; Morgan, R. T.; Zhang, X.; Jacobs, L. M.; Butler, S. G.; Mongora, M. V.; Choy, J. S.; Chen, Y., et al. Naturally occurring mutations of SARS-CoV-2 main protease confer drug resistance to nirmatrelvir. *bioRxiv* **2022**,
